# Forward Programming Identifies Inducers of Blood-Brain Barrier Properties in Human Pluripotent Stem Cell-Derived Endothelial Cells

**DOI:** 10.64898/2026.02.02.699492

**Authors:** Soniya Tamhankar, Yunfeng Ding, Fatemeh Yaghoobi Hashjin, Sarah M. Boutom, Richard Daneman, Sean P. Palecek, Eric V. Shusta

## Abstract

Brain microvascular endothelial cells (BMECs) forming the blood-brain barrier (BBB) maintain brain homeostasis through specialized properties such as tight junctions, efflux transporters, and low levels of transcytosis. However, mechanisms governing induction of BBB properties during development remain poorly understood. We mined single-cell RNA sequencing datasets to identify transcription factors (TFs) critical for BBB development. Forty-four TFs were overexpressed in human pluripotent stem cell-derived endothelial cells cultured in the presence of the Wnt pathway agonist CHIR99021 to identify TFs capable of directing acquisition of BBB properties via forward programming. Individual TFs, including *KLF2*, *KLF4*, *FOXF1*, *FOXF2*, *ZIC2*, *ZIC3*, *NR4A1*, *NR4A2*, *FOXC1*, and *FOXQ1*, induced distinct BBB-like gene expression patterns. Combinations of these TFs induced many canonical BBB genes, yielding ECs with reduced endocytosis, increased efflux activity, and improved barrier function. The resultant forward programmed CNS-like ECs (fpCECs) offer promising tools for modeling human BBB development and neurovascular disease and for drug screening.

## Introduction

The blood-brain barrier (BBB) is a physiological barrier consisting of specialized endothelial cells (ECs) that form the walls of brain blood vessels (*1*) and separate the brain parenchyma from the systemic vasculature. These brain microvascular endothelial cells (BMECs) tightly control the movement of molecules, ions, and cells between the blood and the central nervous system (CNS). They form a tight paracellular barrier via expression of junctional proteins like occludin (OCLN) and claudin-5 (CLDN5) (*2*, *3*), lack fenestrae, and show reduced transcytosis compared to peripheral endothelial cells (*3*, *4*). BMECs further limit non-specific molecular transport through expression of ATP-binding cassette (ABC) efflux transporters, including P-glycoprotein (ABCB1), BCRP (ABCG2), and members of the multidrug resistance-associated protein (MRP) family (*5*). In addition, selective solute uptake into the CNS is mediated by solute carriers (SLCs), such as the neutral amino acid transporter SLC38A5 and glucose transporter GLUT1 (SLC2A1) (*6*). Together, these specialized properties allow BMECs to tightly regulate CNS homeostasis for proper brain function (*1*, *7*, *8*).

The onset of BBB properties takes place during embryonic development as a microenvironment of neural stem cells, pericytes, and neurons provides cues that induce tight junctions and regulate molecular permeability in developing endothelial cells (*9*). In adults, BBB maintenance is a result of composite interactions between brain endothelial cells and resident neural cell types including astrocytes, pericytes, and neurons (*10*). Signaling pathways including Wnt/β-catenin (*11*), retinoic acid (*12*) and netrin signaling (*13*) have been implicated as important actuators of BBB development, in part by regulating BBB-specific transcriptional networks (*14–16*). However, disruption of these pathways reduces, but does not eliminate BBB character (*17*, *18*). Thus, the transcriptional mechanisms responsible for embryonic BBB development and adult BBB maintenance have not been fully elucidated. Transcription factors (TFs) are the master regulators of gene expression, playing a pivotal role in controlling gene expression programs, influencing developmental trajectory and promoting cell lineage-specific characteristics, thereby helping to establish the unique identity of cells (*19*). In particular, other than TFs that act directly downstream of Wnt signaling (e.g. TCF7, LEF1) and retinoic acid receptors, which are likely involved in tuning the Wnt signaling response (*20*, *21*), very little is known about specific TFs that impact BBB formation by orchestrating BBB transcriptional programs.

One valuable tool for identifying TFs capable of inducing BBB properties, particularly in humans, is the use of naïve ECs derived from human pluripotent stem cells (hPSCs). A variety of protocols have been reported to differentiate hPSCs to naïve ECs that lack any demonstrable BBB properties (*22–26*). These ECs can therefore be used to interrogate the capacity of signaling pathways and TFs to induce BBB properties in a developmentally relevant human model. For example, Wnt signaling induction in naïve hPSC-derived ECs was shown to increase expression of GLUT-1 and claudin-5, while decreasing PLVAP expression and permeability, generating CNS-like ECs (CECs) (*27*). In addition, TF overexpression has been used in an attempt to convert non-barrier forming ECs to brain-like ECs. Overexpression of *SOX18*, *SOX7*, *ETS1*, and *LEF1* in hPSC-derived ECs led to modest improvements in barrier as measured by TEER (*28*). Also, the overexpression of *FOXF2* and *ZIC3* individually in human umbilical vein endothelial cells (HUVECs) increased the expression of some BMEC-relevant genes such as *ABCB1*, *SLCO2B1* and *TNFRSF19* (*29*). Although these studies illustrate the potential promise of using hPSC-derived ECs or other EC sources to identify BBB-inducing cues, the resultant cells still lack many BBB properties. Thus, efforts are needed to more comprehensively identify the TFs responsible for imparting BBB identity in developing ECs.

In this work, we explored the ability of TFs that have selectively enriched expression in BMECs for their capacity to induce BBB properties in hPSC-derived CECs (*27*). To this end, BBB-enriched TFs were introduced into hPSC-derived endothelial progenitor cells (EPCs) in concert with Wnt signaling activation. The resultant hPSC-derived CECs transduced with TFs, referred to here as forward programmed CECs (fpCECs), were subjected to RNA sequencing, molecular phenotyping, and functional analyses to identify a subset of TFs capable of inducing a range of BBB properties. Moreover, synergistic combinations of TFs were identified that further drive BBB specification in fpCECs.

## Results

### Mining single cell transcriptomics data to identify BBB-enriched transcription factors

To identify TFs with enriched expression in BBB endothelial cells, we performed independent analyses of single cell transcriptomics datasets. In the first analysis, we compared the transcriptomes of adult murine brain EC populations with murine ECs from five peripheral vascular beds: heart, lung, liver, kidney and skeletal muscle (*30*), generating a list of 48 TFs that were enriched in murine brain EC populations (Fig. 1). For the second analysis, we pooled adult human brain ECs from five different single cell transcriptomics datasets (*31–35*), and compared brain EC gene expression to pooled human heart, liver, lung and skeletal muscle ECs (*36–39*) to identify 55 TFs with enriched expression in human brain EC populations (Fig. 1) (*40*). Finally, from an embryonic human brain single cell transcriptomics dataset, we compared non-barrier forming brain tip cells with brain capillary endothelial cells to identify 40 TFs enriched in those cells that acquired a transcriptomic BBB signature (*41*). From this integrated analysis, we identified 40 TFs that exhibited increased expression in brain ECs during or after barriergenesis in at least two out of the three datasets, and 11 TFs were enriched in all three comparisons (Fig. 1).

**Fig. 1.**
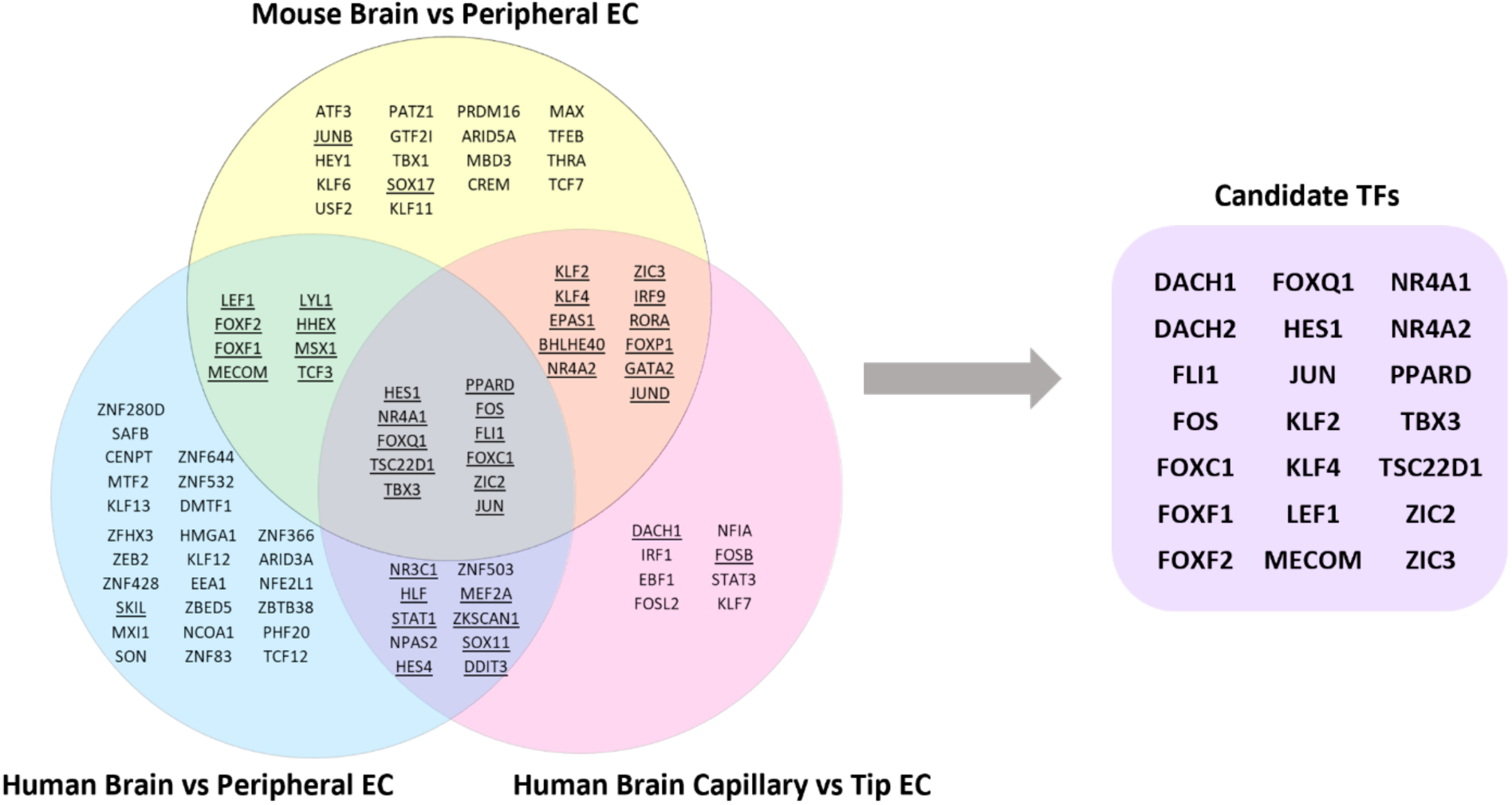
Single cell transcriptomic analyses reveal BBB-enriched TFs. Publicly available single cell transcriptomic datasets were analyzed to determine BBB-enriched TFs (Non-parametric Wilcoxon rank sum test, log_2_ fold change>1, and p-value < 0.05) (*27*, *30–39*, *41*). A total of 92 TFs were identified as shown in the Venn diagram. 40 TFs were significantly enriched in the brain EC or brain capillary EC populations in at least two of the three independent transcriptomics comparisons. Using a preliminary qPCR screen for 44 TFs (underlined) and detailed in Supplementary Figure 1, we determined a list of 21 candidate TFs for individual overexpression and bulk transcriptomic analysis in hPSC-derived CNS-like ECs.

To narrow the list of candidate TFs, we transduced hPSC-derived CECs with individual TFs and assessed their effects on a panel of BBB transcripts. The hPSC-derived CECs were chosen as the cellular platform since they possess partial BBB character as a result of Wnt signaling activation in hPSC-derived EPCs (*27*) and could therefore be used to identify TFs that could further increase BBB properties. To this end, hPSCs were differentiated through mesoderm progenitors to EPCs as previously described (*27*, *42*). After EPC purification by CD31+ magnetic-activated cell sorting (MACS), lentiviruses delivering individual TFs were dosed onto EPCs with simultaneous treatment of Wnt agonist CHIR99021, yielding fpCECs (Fig. 2A). From the list of 40 TFs enriched in at least 2 of the 3 analyses, we excluded *ZNF503* and *NPAS2* from screening due to availability. Six additional TFs, five of which were enriched in 1 out of the 3 datasets (*SOX17, FOSB, JUNB, SKIL*, *DACH1*) along with *DACH2* were included. *SOX17* was included due to previously reported role in regulating BBB properties (*43*). *SKIL* (*44*), and *DACH1* with its paralog *DACH2* were included due to their known effects on regulating TGF-β signaling (*45*). *FOSB* and *JUNB* were included because they, along with *FOS* and *JUND*, are crucial members of the AP-1 signaling cascade (*46*). Resultant fpCECs harboring single TFs or control lentivirus were evaluated by RT-qPCR for a panel of 8 genes enriched at the BBB (*CLDN5*, *OCLN*, *ABCB1*, *ABCG2*, *SLC2A1*, *SLC7A5*, *SLC38A5*, *MFSD2A*), 2 genes downregulated at the BBB (*CAV1*, *PLVAP*), and 2 endothelial cell markers (*CDH5*, *PECAM1*) (fig. S1). We found that out of the 44 tested TFs, many induced some transcriptional aspects of BBB character. For example, *KLF2* and *KLF4* increased the expression of *MFSD2A*; *ZIC2*, *ZIC3*, *KLF2* and *KLF4* increased the expression of *SLC38A5;* and *KLF2* increased the expression of *ABCB1* (fig. S1). We then selected TFs that either induced BBB gene expression in hPSC-derived CECs in the RT-qPCR assay or TFs that were enriched in all three datasets to narrow the list of TFs to 21 candidates for comprehensive transcriptomic characterization (Fig. 1).

**Fig. 2.**
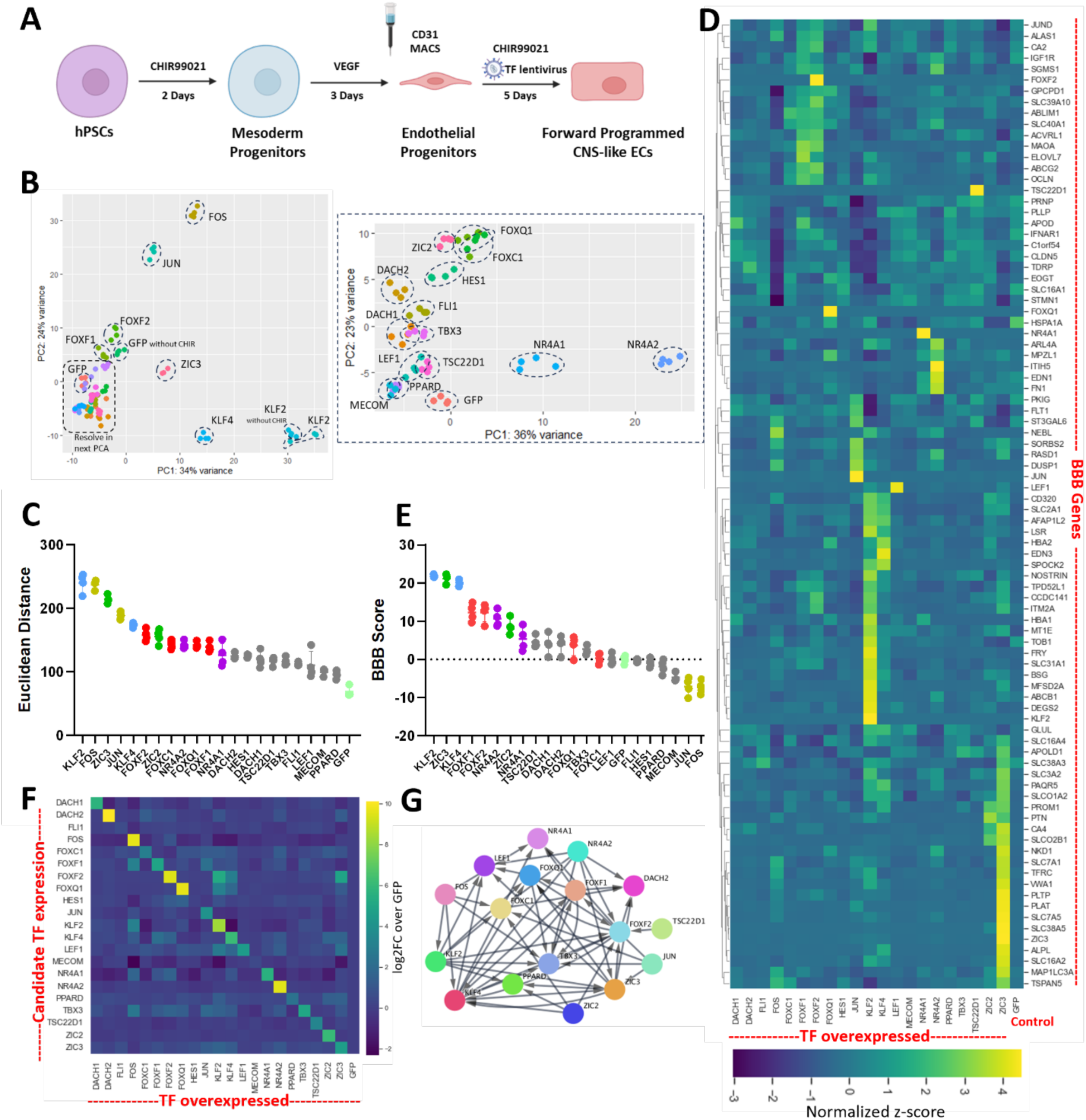
fpCECs with single TF overexpression have improved BBB-like gene expression. **(A)** IMR90-4 hPSCs were differentiated to endothelial progenitor cells (EPCs). CD31+ EPCs were MACS-sorted. Lentivirus for TF overexpression was dosed on CD31+ EPCs and the population was simultaneously treated with CHIR99021 to activate Wnt signaling. After five days, the resultant cell population is referred to as forward programmed CNS-like ECs (fpCECs). Bulk RNA sequencing was performed on the fpCECs at N=4 independent lentivirus transductions for each condition. **(B)** Principal component analysis (PCA) was performed on fpCECs and CECs overexpressing GFP as a control. A second PCA (right) was performed on samples within the dashed box in the first PCA (left) to better resolve the more closely related TF-driven transcriptomes. Each sample on the graphs represents an independently sequenced biological replicate. **(C)** Euclidean distance between the transcriptomes of fpCECs and the average transcriptome of CECs overexpressing GFP is plotted (see Methods for details). The mean±S.D. of N=4 sequencing replicates is plotted. **(D)** Heatmap of the expression of 88 BBB-enriched genes for fpCECs overexpressing each of the 21 candidate TFs. Each row represents z-scores normalized to the mean expression for each of the 88 BBB-enriched genes. The rightmost column is the GFP control. Average z-score of N=4 biological replicates is plotted. **(E)** BBB score analysis of fpCECs overexpressing each of the 21 candidate TFs. See the Methods for BBB score calculation details. The mean±S.D. of N=4 sequencing replicates is plotted. Colors for individual TFs in plots (C) and (E) match. **(F)** Heatmap of the expression of each of the 21 candidate TFs upon overexpression of each individual TF on the x-axis. Average Log_2_ fold-changes compared to the GFP control are plotted for N=4 biological replicates. **(G)** String-style map for gene expression regulation amongst the 21 candidate TFs. For example, if TF A overexpression drives the upregulation of TF B more than two-fold in a statistically significant manner, an arrow from TF A to TF B is plotted. A p_adj_<0.05 using the Wald test followed by Benjamini-Hochberg correction was used as the statistical significance cutoff. TFs not represented on the string map did not have their expression affected (> 2-fold) by any other overexpressed TF. Full transcriptome expression data for all TFs and their replicates can be found in Supplementary File 1)

### BBB-enriched TFs induce BBB-like gene expression in fpCECs

Given that the qPCR screen identified TFs that induced expression of a subset of BBB-related genes, we assessed the global transcriptomic changes elicited by overexpression of individual TFs via bulk RNA sequencing analysis on fpCECs individually transduced with each of the 21 TFs (Fig. 2A). CECs transduced with a GFP lentivirus were used as controls. Cells from a parallel transduced sample were used for quality control to ensure that transduced cells remained pure CDH5-expressing EC populations (fig. S2). A principal component analysis (PCA) of all sequenced transcriptomic profiles revealed that transcriptional changes induced by overexpression of *KLF2* and *KLF4* TFs dominated PC1, while changes induced by overexpression of AP-1 family TFs *FOS* and *JUN* dominated PC2 (Fig. 2B). *FOXF1*, *FOXF2* and *ZIC3* also induced significant changes in gene expression (Fig. 2B). Euclidean distances between transcriptomic profiles to the average of GFP controls demonstrated that *KLF2*, *FOS*, *ZIC3*, *JUN*, *KLF4* and *FOXF2* induced the most significant global gene expression changes (Fig. 2C). To specifically evaluate the impact of TF overexpression on BBB specification, we curated a list of 88 genes that were upregulated in murine brain ECs compared to peripheral ECs in a single cell sequencing study (*30*), while still having significant expression levels in human brain EC datasets (Table S1, details in Methods). We found that although no single TF induced all 88 BBB genes, *KLF2* or *KLF4*, *FOXF1* or *FOXF2*, *ZIC3*, *NR4A1* or *NR4A2*, and *FOXQ1* each induced fairly discrete BBB gene subsets (Fig. 2D, file S1 and S2). To quantify these changes, we calculated a BBB score to measure the collective extent of TF overexpression on the induction of the 88 gene BBB transcriptional program (details in Methods) and identified that pairs of homologs *KLF2* and *KLF4*, *ZIC2* and *ZIC3*, *FOXF1* and *FOXF2*, *NR4A2* and *NR4A1* had the highest BBB scores, indicating their capability to induce BBB-like gene expression changes in fpCECs (Fig. 2E). We then probed whether any of the 21 TFs can regulate the expression of other BBB-enriched TFs and identified significant crosstalk (Fig. 2F, file S2). For example, overexpression of *KLF2* induced the expression of *LEF1*, *FOXF2*, *FOXQ1* and *ZIC3* while overexpression of *ZIC3* induced the expression of *FOXF2*, *FOXQ1*, *KLF4* and *LEF1*. These relationships were also summarized in a STRING plot (Fig. 2G), which identified KLF family TFs, ZIC family TFs, FOX family TFs and NR4A family TFs to be the TFs most capable of increasing the expression of other BBB-enriched TFs.

### *KLF2* and *KLF4* induce BBB-like gene and protein expression in fpCECs

*KLF2* and *KLF4*, zinc finger proteins belonging to the Krüppel-like factors family of TFs, play important roles in endothelial cells. For example, shear stress induces MEK5/ERK5 signaling, which in turn increases *KLF2* and *KLF4* expression. KLF family TF expression regulates endothelial gene expression, increases chromatin accessibility, regulates vascular inflammation and induces vasoprotective effects (*47*, *48*). In single cell transcriptomics datasets, *Klf2* and *Klf4* were upregulated in murine brain endothelial cells compared to non-brain endothelial cells (Fig. 3, A and B) (*30*). In addition, *KLF2* and *KLF4* were upregulated in human brain capillary endothelial cells compared to tip cells (Fig. 3, C and D) (*41*). When analyzing the effect of *KLF2* or *KLF4* expression in fpCECs, we found that *KLF2* increased the expression of 46 of the 88 BBB-enriched genes and decreased the expression of 30 of these genes. Similarly, *KLF4* increased the expression of 36 and decreased the expression of 25 of the 88 BBB-enriched genes. (Fig. 3F, file S2). Specifically, *KLF2* or *KLF4* overexpression increased the expression of BBB-enriched TFs, including *FOXF2*, *FOXQ1* and *LEF1* at relatively high fold changes, indicating that *KLF2* or *KLF4* may serve as upstream regulators (Fig. 3E, fig. S3A). *KLF2* or *KLF4* overexpression also significantly induced the expression of efflux transporter transcripts *ABCB1* and *ABCG2*, specialized transporter or transporter-related transcripts *MFSD2A*, *SLC16A2*, *SLC2A1*, *SLC31A1*, *SLC38A5*, *SLC7A5*, *SLCO1A2*, *SLCO2B1* and *TFRC*, and adhesion and junction-related genes *BSG*, *LSR* and *OCLN* (Fig. 3G, file S2). *KLF2* or *KLF4* overexpression also increased the expression of several BBB-enriched enzyme and signaling-related transcripts (fig. S3B, file S2). *KLF2* or *KLF4* overexpression reduced the expression of *CAV1* and *CAV2* transcripts, key molecular components of caveolae-mediated endocytosis (Fig. 3E). We validated transcript-level changes by probing a subset of protein targets via immunocytochemistry (ICC), and found that on a protein level, *KLF2* overexpression significantly increased the expression of ABCB1, ABCG2, LSR, SLC2A1, SLC38A4 and OCLN, while significantly decreasing the expression of CAV1 (Fig. 3, H and I). Similarly, *KLF4* overexpression increased protein expression of LEF1, LSR, SLCO2B1, SLC38A5, OCLN, and decreased expression of CAV1 (fig. S3, C and D). While several proteins were upregulated in all transduced cells, others like OCLN were upregulated in a subset of transduced cells. Since it has been suggested that *KLF2* and *KLF4* are targets of ERK5 signaling, we analyzed the expression of known genes downstream of ERK5 signaling in endothelial cells (*49*), and found that *KLF2* or *KLF4* overexpression significantly increased the expression of such genes (Fig. 3J, file S2).

**Fig. 3.**
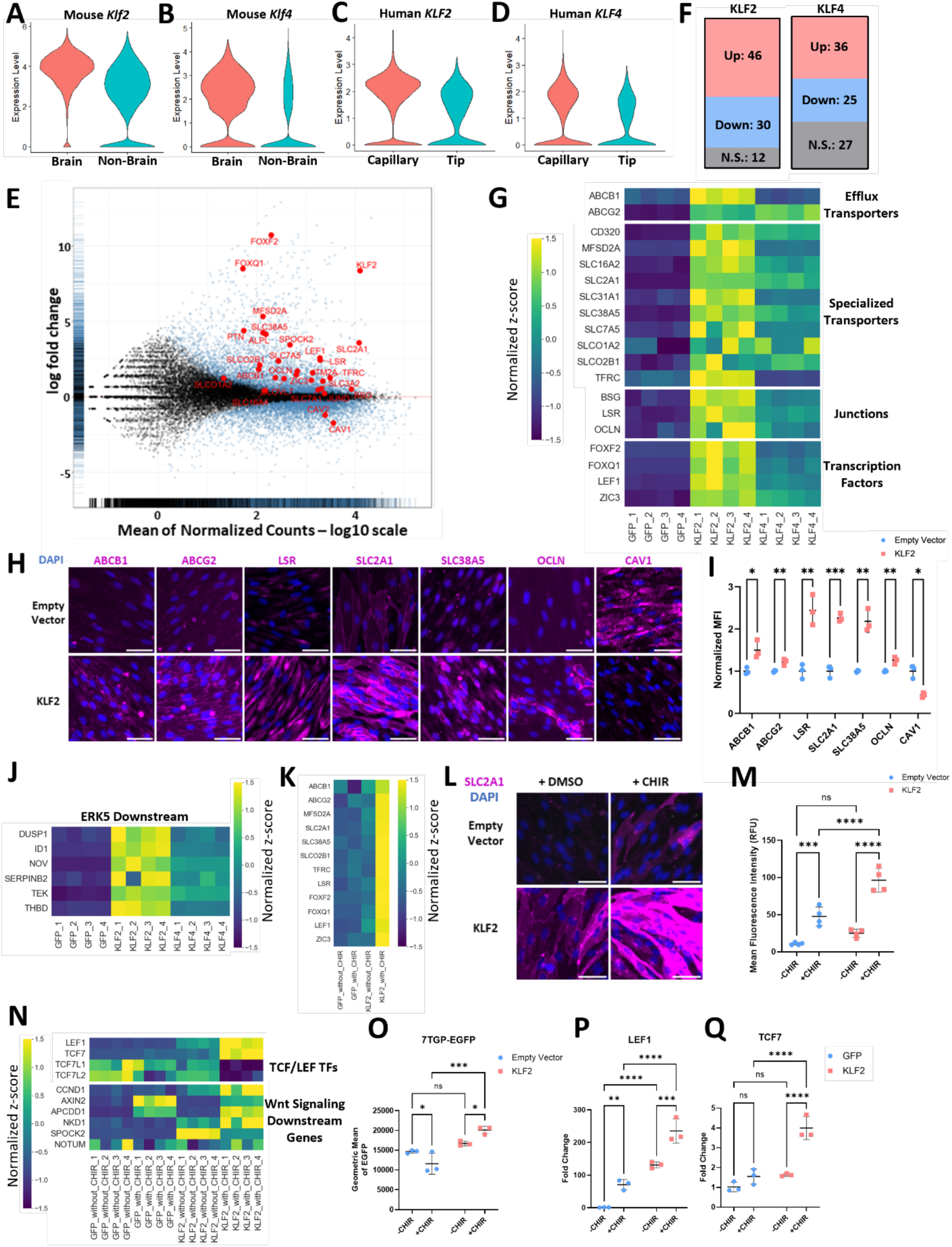
KLF family TFs induce significant BBB-like gene and protein expression through synergy with canonical Wnt signaling. Expression of **(A)** *Klf2* and **(B)** *Klf4* in mouse brain and non-brain (heart, lung, liver, kidney and skeletal muscle) ECs from a single cell transcriptomic analysis (*30*). Expression of **(C)** *KLF2* and **(D)** *KLF4* in human brain capillary and tip ECs (*41*). **(E)** Log ratio-average (MA) plot for differential gene expression between fpCECs overexpressing *KLF2* and CECs overexpressing GFP. X-axis is the average log_10_ of normalized counts in RNA-seq. Y-axis is the log_2_ fold change of gene expression. Select BBB-enriched and depleted genes are marked red. **(F)** Number of the 88 BBB-enriched genes upregulated, downregulated, or exhibiting no significant changes upon overexpression of *KLF2* or *KLF4* in fpCECs. Adjusted p-value < 0.05 using the Wald test was used to determine whether upregulation or downregulation was statistically significant. **(G)** Heatmap of the expression of efflux transporters, specialized transporters, junction genes and transcription factors that were upregulated by *KLF2* and *KLF4* overexpression. Each row represents z-scores normalized to the mean expression for each gene. N=4 biological replicates per condition. fpCECs received either a control GFP lentivirus, *KLF2* lentivirus, or *KLF4* lentivirus at MOI=2. **(H)** Immunocytochemistry of efflux transporter (ABCB1, ABCG2), specialized transporter (SLC2A1, SLC38A5), junction (LSR, OCLN) and endocytosis-related proteins (CAV1) in fpCECs receiving either a control empty vector lentivirus, or *KLF2* lentivirus at MOI=2. Scale bar is 50 µm. **(I)** Quantification of immunocytochemistry images. N=3 biological replicates per condition. Signals were normalized to the average of mean fluorescence intensities (MFIs) in samples receiving empty vector virus for each protein assessed. *:p<0.05; **:p<0.01; ***:p<0.001 in Student’s t-tests. **(J)** Heatmap of the expression of target genes downstream of the ERK5 signaling pathway. N=4 biological replicates per condition. fpCECs received either a control GFP lentivirus, *KLF2* lentivirus, or *KLF4* lentivirus at MOI=2. **(K)** Heatmap of the expression of a panel of BBB enriched genes with functional annotation, including efflux transporters (*ABCB1, ABCG2*), specialized transporters (*MFSD2A, SLC2A1, SLC38A5, SLCO2B1, TFRC*), tight junction protein (*LSR*), and transcription factors (*FOXF2, FOXQ1, LEF1, ZIC3*). Samples are EPCs receiving either Wnt agonist CHIR99021 or DMSO control, and *KLF2* lentivirus or GFP control lentivirus. Average of N=4 biological replicates per condition are plotted**. (L)** Immunocytochemistry of glucose transporter SLC2A1in EPCs receiving either Wnt agonist CHIR99021 or DMSO control, and *KLF2* lentivirus or empty lentivirus. Scale bar is 50µm. **(M)** Quantification of SLC2A1 immunocytochemistry images. N=4 biological replicates per condition. ***:p<0.001; ****:p<0.0001 in two-way ANOVA followed by Tukey’s post-hoc HSD test. **(N)** Heatmap of the expression of canonical Wnt signaling TCF/LEF family TFs, and downstream genes of Wnt signaling. Samples are EPCs receiving either Wnt agonist CHIR99021 or DMSO control, and *KLF2* lentivirus or GFP control lentivirus. Average of N=4 biological replicates per condition are plotted. **(O)** Geometric mean of eGFP expression using 7xTCF-eGFP H9 (7TGP) Wnt reporter line-derived EPCs transduced with either empty vector or *KLF2* lentivirus and cultured with or without CHIR99021. Flow data can be found in Figure S4F. N=3 biological replicates per condition. *:p<0.05; ***:p<0.001 in two-way ANOVA followed by Tukey’s post-hoc HSD test. **(P-Q)** qPCR quantification of *LEF1*, *TCF7*, and *SLC2A1* gene expression in HUVECs transduced with GFP or *KLF2* lentivirus and cultured with or without CHIR99021. Expression is normalized to samples treated with empty vector lentivirus in the absence of CHIR99021. N=3 biological replicates, **:p<0.01; ***:p<0.001, ****:p<0.0001 in two-way ANOVA followed by Tukey’s post-hoc HSD test.

In the hPSC-derived fpCECs, Wnt signaling was activated via the GSK3β agonist CHIR99021 concordant with TF transduction and overexpression (Fig. 2A). Recognizing the strong effect of *KLF2* or *KLF4* overexpression on BBB gene expression, we explored whether activated Wnt signaling impacted the ability of KLF overexpression to modulate BBB gene expression. We sequenced both GFP-transduced control and *KLF2* overexpression samples with or without CHIR99021 exposure. Several key BBB genes, including *ABCB1*, *ABCG2*, *MFSD2A*, *SLC2A1*, *SLC38A5*, *SLCO2B1*, *TFRC*, *LSR*, *FOXF2*, *FOXQ1*, *LEF1* and *ZIC3* were only upregulated in samples having both *KLF2* overexpression and Wnt activation by CHIR99021 (Fig. 3K, fig. S4A, file S2). This finding indicates a synergistic effect of these two pathways in inducing BBB gene expression. Further validation at the protein level indicated that the induction of protein expression of SLC2A1, SLC38A5 and SLCO2B1 by *KLF2* overexpression, required Wnt co-activation (Fig. 3, L and M, fig. S4, B-E). Since several of the genes synergistically induced by *KLF2* overexpression and CHIR99021 treatment are also known targets of canonical Wnt signaling, such as *SLC2A1, LEF1* and *SLC38A5* (*20*), we analyzed the expression of several key TCF/LEF TFs and downstream targets. We found that combining *KLF2* overexpression with CHIR99021 treatment significantly increased the expression of canonical Wnt signaling activators *LEF1/TCF7* and reduced the expression of Wnt inhibitory TFs *TCF7L1* and *TCF7L2* (Fig. 3N). We also observed upregulation of *CCND1*, a canonical downstream target of TCF/LEF signaling, as well as increased expression of Wnt signaling downstream targets (*AXIN2*, *APCDD1*, *NKD1*, *SPOCK2* and *NOTUM*) previously identified in a *Ctnnb1* mouse endothelial knockout model (*18*) (Fig. 3N, file S2). Such shifts in Wnt TF expression have been reported to be correlated with β-catenin levels, and represent higher levels of canonical Wnt activity. To further assess the canonical Wnt signaling activation, we used an hPSC Wnt reporter cell line, H9 7TGP (*50*, *51*). After differentiating H9 7TGP hPSCs to EPCs (Fig. 2A), they were treated with *KLF2* lentivirus and/or CHIR99021. We found that while *KLF2* overexpression alone did not induce GFP expression in H9 7TGP fpCECs, *KLF2* overexpression in conjunction with CHIR99021 treatment significantly induced GFP expression over CHIR99021 treatment alone (Fig. 3O, fig. S4F), indicating stronger canonical Wnt signaling activity with combined *KLF2* and CHIR99021 treatment. To assess whether *KLF2* and CHIR99021 treatment also induces BBB gene expression in primary ECs, we treated HUVECs with *KLF2* lentivirus and CHIR99021. While either *KLF2* or CHIR99021 treatment significantly increased the expression of canonical Wnt signaling TF transcripts *LEF1* and *TCF7* (Fig. 3, P-Q), the expression of BBB markers *SLC2A1* and *OCLN* were only upregulated in the *KLF2*-treated HUVECs (fig. S4G).

Taken together, *KLF2* and *KLF4* TFs induced significant BBB-like gene and protein expression. Interestingly, these effects were significantly more pronounced when *KLF2* or *KLF4* overexpression was accompanied by Wnt activation. In addition, the expression of key BBB-enriched TFs, including *FOXF2*, *FOXQ1* and *ZIC3*, were only significantly upregulated when *KLF2* or *KLF4* overexpression and Wnt activation were both present.

### FOX, ZIC and NR4A family transcription factors induce BBB-like gene expression

Overexpression of Forkhead box (Fox) TFs *FOXF1*, *FOXF2*, *FOXQ1*, and *FOXC1* also improved BBB scores in fpCECs (Fig. 2E). Notably, expression of these four TFs is almost absent in non-brain EC populations in mice and humans, indicating their BBB vascular specificity (Fig. 4, A-D, I-L) (*30*, *41*). When analyzing the specific effects of *FOXF1* or *FOXF2* on BBB-enriched genes in fpCECs, we found that *FOXF1* or *FOXF2* overexpression increased the expression of BBB-enriched TFs *FOXQ1* and *NR4A1*, suggesting synergistic regulation amongst these TFs (Fig. 4, E-F, file S2). We also found that *FOXF1* or *FOXF2* overexpression significantly induced the expression of efflux transporter *ABCG2*, specialized nutrient transporters *SLC2A1*, *SLC31A1*, *SLC39A10*, *SLC40A1*, and *TFRC,* junctional gene *OCLN,* and genes involved in lipid metabolism (Fig. 4F, fig. S5, A and B, file S2). In addition, both *FOXF1* and *FOXF2* induced SLC2A1 (GLUT-1) protein expression (Fig. 4, G and H).

**Fig. 4.**
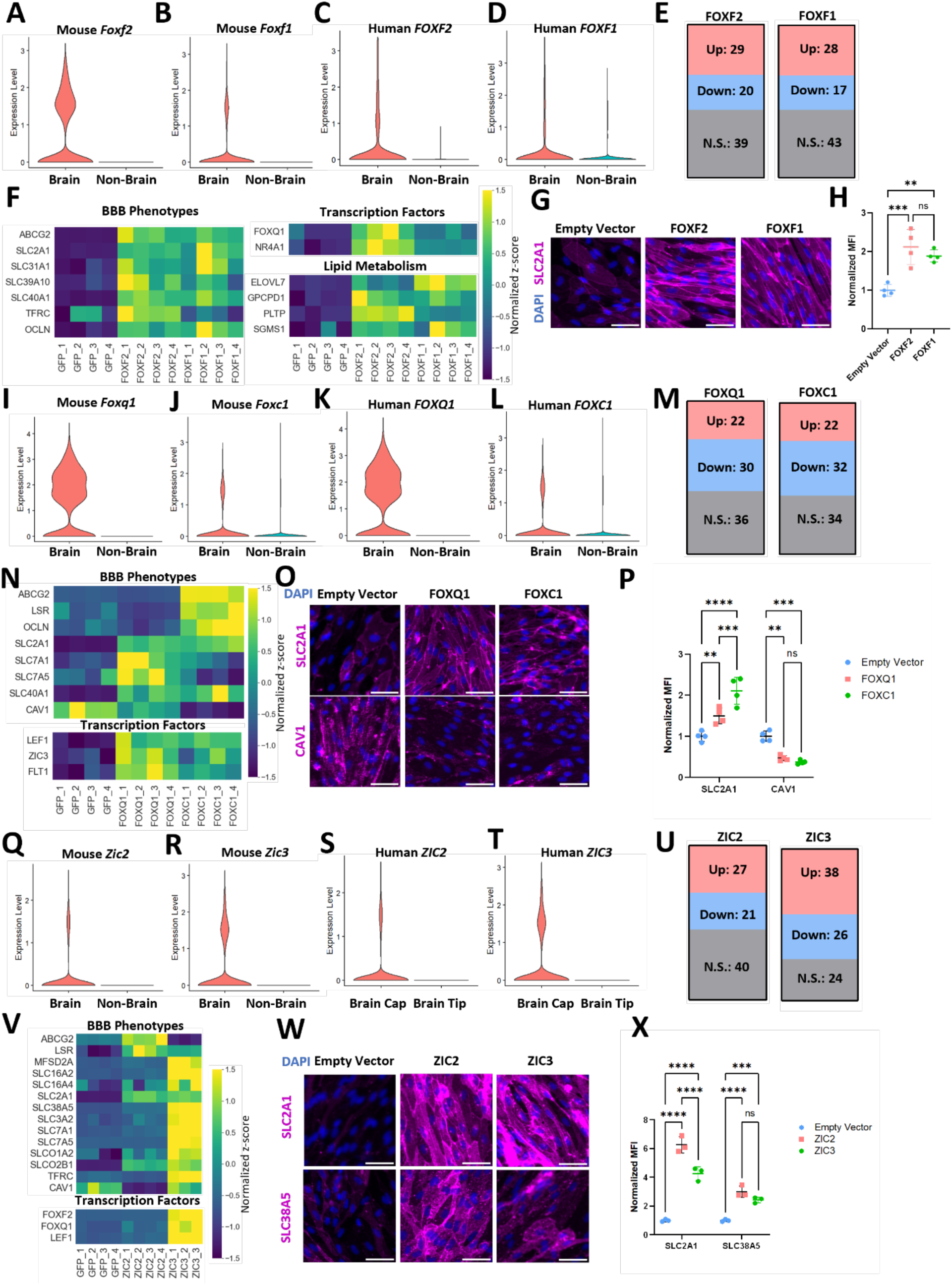
Figure 4. FOX and ZIC family TFs induce significant BBB-like gene and protein expression. Expression of **(A)** *Foxf2* and **(B)** *Foxf1* in mouse brain and non-brain (heart, lung, liver, kidney and skeletal muscle) ECs in single cell transcriptomic analysis (*30*). Similarly, expression of **(C)** *FOXF2* and **(D)** *FOXF1* in human brain ECs and non-brain (heart, liver, lung and skeletal muscle) ECs was plotted (*27*, *31–39*). **(E)** Number of 88 BBB-enriched genes upregulated, downregulated, or exhibiting no significant changes upon overexpression of *FOXF1* or *FOXF2* in fpCECs. Adjusted p value < 0.05 using the Wald test was used to determine whether upregulations and downregulations are statistically significant. **(F)** Heatmap of the expression of genes related to BBB phenotypes, transcription factors and lipid metabolism upregulated by *FOXF2* and *FOXF1* overexpression in fpCECs. z-scores were normalized to the mean expression for each of the BBB-enriched genes. N=4 biological replicates per condition. fpCECs received either a control GFP or TF lentivirus at MOI=2. **(G)** Immunocytochemistry of SLC2A1 in fpCECs receiving either an empty vector lentivirus, or TF lentivirus at MOI=2. Scale bar is 50µm. **(H)** Quantification of SLC2A1 immunocytochemistry images. N=4 biological replicates per condition. MFI: mean fluorescence intensity. **:p<0.01, ***:p<0.001 in one-way ANOVA followed by Tukey’s post-hoc HSD test. Expression of **(I)** *Foxq1* and **(J)** *Foxc1* in mouse brain and non-brain (heart, lung, liver, kidney and skeletal muscle) ECs in single cell transcriptomic analysis. Similarly, expression of **(K)** *FOXQ1* and **(L)** *FOXC1* in human brain ECs and non-brain (heart, liver, lung and skeletal muscle) ECs was plotted. **(M)** Number of 88 BBB-enriched genes upregulated, downregulated, or exhibiting no significant changes upon overexpression of *FOXQ1* or *FOXC1* in fpCECs. Adjusted p value < 0.05 using the Wald test was used to determine whether upregulation or downregulation was statistically significant. **(N)** Heatmap of the expression of genes related to BBB phenotypes and transcription factors that are upregulated by *FOXQ1* and *FOXC1* overexpression. Z-scores normalized to the mean expression for each of the BBB-enriched genes are plotted. N=4 biological replicates per condition. fpCECs receiving either a control GFP lentivirus, or TF lentivirus at MOI=2. **(O)** Immunocytochemistry of SLC2A1 and CAV1 in fpCECs receiving either an empty vector lentivirus, or TF lentivirus at MOI=2. Scale bar is 50µm. **(P)** Quantification of immunocytochemistry images. N=4 biological replicates per condition. MFI: mean fluorescence intensity. **: p<0.01; ***:p<0.001; ****:p<0.0001 in one-way ANOVA followed by Tukey’s post-hoc HSD test. Expression of **(Q)** *Zic2* and **(R)** *Zic3* in mouse brain and non-brain (heart, lung, liver, kidney and skeletal muscle) ECs in single cell transcriptomic analysis. Similarly, expression of **(S)** *ZIC2* and **(T)** *ZIC3* in human brain capillary and tip ECs was plotted. **(U)** Number of 88 BBB-enriched genes upregulated, downregulated, or exhibiting no significant changes upon overexpression of *ZIC2* or *ZIC3* in fpCECs. Adjusted p value < 0.05 using the Wald test was used to determine whether upregulation or downregulation was statistically significant. **(V)** Heatmap of the expression of genes related to BBB phenotypes and transcription factors that are upregulated by ZIC family TF overexpression. z-scores normalized to the mean expression for each of the BBB-enriched genes are plotted. N=4 biological replicates per condition. fpCECs receiving either a control GFP lentivirus, or TF lentivirus at MOI=2. **(W)** Immunocytochemistry of SLC2A1 and SLC38A5 in fpCECs receiving either an empty vector lentivirus, or TF lentivirus at MOI=2. Scale bar is 50µm. **(X)** Quantification of immunocytochemistry images. N=4 biological replicates per condition. MFI: mean fluorescence intensity. ***:p<0.001; ****:p<0.0001 in one-way ANOVA followed by Tukey’s post-hoc HSD test.

From PCA analysis, overexpression of *FOXC1* and *FOXQ1* generated similar transcriptomic profiles (Fig. 2B). *FOXC1* or *FOXQ1* overexpression in fpCECs increased the expression of BBB-enriched TFs *LEF1*, *ZIC3* and *FLT1* (Fig. 4, M and N, file S2). *FOXC1* overexpression significantly induced the expression of transporter genes *ABCG2* and *SLC2A1* and tight junction genes *LSR* and *OCLN,* while *FOXQ1* overexpression more prominently induced the collective expression of specialized transporters *SLC2A1*, *SLC7A1*, *SLC7A5* and *SLC40A1* (Fig. 4N, fig. S5, C and D). Both *FOXC1* and *FOXQ1* reduced the expression of *CAV1*. Concordant with the transcript level changes, protein expression of SLC2A1 was significantly increased and CAV1 was significantly decreased by *FOXQ1* and *FOXC1* overexpression (Fig. 4, O and P).

Overexpression of *ZIC2* and *ZIC3,* TFs that play important roles in pluripotency and nervous system development (*52*), also significantly enhanced the BBB score in fpCECs (Fig. 2E). *ZIC2* and *ZIC3* transcripts are almost absent in mouse and human non-brain EC populations, indicating specificity for the BBB (Fig. 4, Q-T). *ZIC3* overexpression induced more significant BBB-like transcriptomes than *ZIC2* in fpCECs (Fig. 2E, Fig. 4, U and V, file S2, fig. S5, E and F). Notably, *ZIC3* overexpression increased the expression of other BBB-enriched TFs including *FOXF2*, *FOXQ1,* and *LEF1* (Fig. 4V, file S2). *ZIC3* overexpression also significantly induced the expression of BBB-enriched genes *MFSD2A*, *SLC16A2*, *SLC16A4*, *SLC2A1*, *SLC38A5*, *SLC3A2*, *SLC7A1*, *SLC7A5*, *SLCO1A2*, *SLCO2B1* and *TFRC* (Fig. 4V, file S2). Overexpression of either *ZIC2* or *ZIC3* increased protein expression of SLC2A1 and SLC38A5 (Fig. 4, W and X).

Nuclear Receptor subfamily 4, group A (NR4A) family TFs serve as essential regulators for development and maintenance of many neural, immune and muscle cell types (*53*). Overexpression of *NR4A1* or *NR4A2* improved BBB scores in fpCECs (Fig. 2E). Expression of *NR4A1* and *NR4A2* is almost absent in non-brain EC populations in mice, while *NR4A1* and *NR4A2* expression is more abundant in human brain capillary ECs than brain tip ECs (fig. S6, A-D). Overexpression of either TF in fpCECs significantly induced the expression of a myriad of BBB-enriched transporters including *ABCG2*, *SLC2A1*, *SLC38A5*, *SLC7A1*, *SLC7A5*, *SLC40A1* (fig. S6, E-H, file S2), and BBB-enriched TFs LEF1 and *ZIC3* (fig. S6H). Upregulation of SLC7A5 was validated at the protein expression level (fig. S6, I and J). Overall, FOX, ZIC and NR4A family TFs induced significant, distinct BBB-like transcriptional character in fpCECs.

While overexpression of *KLF2*, *KLF4*, *FOXF1*, *FOXF2, FOXQ1*, *FOXC1, ZIC2, ZIC3, NR4A1* and *NR4A2* in fpCECs resulted in pure populations of CDH5+ ECs (fig. S2), we observed reduced transcript expression of endothelial markers *PECAM1* and *CDH5* for some of these TFs such as *KLF2* (fig. S7A), a gene expression pattern also captured in single cell RNA sequencing data sets when comparing mouse brain ECs with peripheral ECs (fig. S7B). However, the observation of reduced expression of *PECAM1* and *CDH5* seems to be restricted to the hPSC-derived endothelial system, as combined treatment of *KLF2* and CHIR99021 did not decrease expression of endothelial markers *PECAM1* and *CDH5* in HUVECs (fig. S7C). When inspecting the gene expression of endothelial (*CDH5, CD34, PECAM1, CLDN5, ERG, FLI1*), mesenchymal (*PDGFRB, CSPG4, PDGFRA, TBX2, CNN1, COL1A1*) and epithelial (*CDH1, EPCAM, CLDN1, CLDN3, CLDN4, CLDN6*) transcripts, we found that fpCECs overexpressing any of these 10 individual TFs still retained an endothelial signature (fig. S7D, file S2).

### Combinatorial expression of BBB-enriched TFs enhances BBB-like gene and protein expression

Realizing that overexpression of single TFs induced distinct subsets of the BBB gene expression program (Fig. 2D), we hypothesized that combining the expression of multiple transcriptional regulators could achieve a broader recapitulation of BBB gene expression. To test this, we first calculated the average transcriptional profile changes in fpCECs induced by the individual overexpression of each of the 21 TFs and then predicted how combinatorial overexpression of 4 or 5 TFs would induce transcriptomic shifts similar to those observed when comparing BBB-like ECs to non-BBB-like ECs (Details in Methods). An averaged transcriptomic profile approach was chosen based on a previous finding that averaging transcriptomics profiles outperforms other methods in predicting effects of combinatorial TF overexpression (*19*). We found that for predicted combinations of four or five TFs, combinations containing *KLF2*, *KLF4*, *FOXF1*, *FOXF2*, *ZIC3*, *NR4A2*, and *FOXQ1* consistently generated the highest prediction scores (file S3). Based on these predictions, we chose 11 TF combinations to test experimentally in fpCECs (Fig. 5A). When compared to BBB gene expression changes upon single TF overexpression (Fig. 2D, file S2), expression of a combination of TFs induced a broader set of BBB genes (Fig. 5B, file S2) compared to GFP control. Multiple TF overexpression was confirmed for each TF combo (Fig. 5C). Principal component analysis delineated the impact of KLF family TF overexpression along the diagonal of PC1 and PC2, and *ZIC3* overexpression along PC2, with each combination leading to transcriptomes distinct from those generated by overexpression of any single TF (Fig. 5D). We again performed BBB score analysis to quantify enrichment of BBB genes and found that all 11 TF combinations tested gave higher BBB scores than any single TF (Fig. 5F), indicating combinatorial BBB-inducing impact for these lead TF candidates. Notably, the TF combinations with top BBB scores (Combos 4, 5, 9) exhibited smaller overall transcriptomic Euclidean distances from GFP control compared to single TF overexpression with *KLF2* (Fig. 5E), suggesting a selective impact of these TFs on BBB gene expression rather than just reflecting a global shift in gene expression. As suggested by the elevated BBB scores, we found broad impacts on the expression of BBB-enriched genes, with 64, 60 and 64 of the 88 genes being increased by combos 4, 5 and 9, respectively (Fig. 5G, file S2). We then assessed the expression of the 40 BBB-enriched TFs shown in Fig. 1 that were not part of the TF overexpression combinations, and found that the top TF combinations (combos 4, 5, 9) induced expression of many of the other BBB-enriched TFs, suggesting that these discrete 4-5 member TF combinations have the potential to initiate a broader BBB-specification transcriptional program (fig. S8A, file S2). When comparing the BBB-enriched gene expression elicited by TF combinations with that generated by overexpressing the individual TFs that comprise the combo, we found that subsets of the BBB-enriched gene expression profiles were shared between individual TF and TF combination samples as expected. There were also many instances where TF combinations resulted in significantly enhanced BBB-enriched gene expression compared with that resulting from individual TFs (fig. S8B, file S2). Moreover, there were also instances where TF combinations reduced the level of induction compared to single TF overexpression. These observations suggest synergistic interplay amongst these TFs in inducing BBB-like gene expression.

**Fig. 5.**
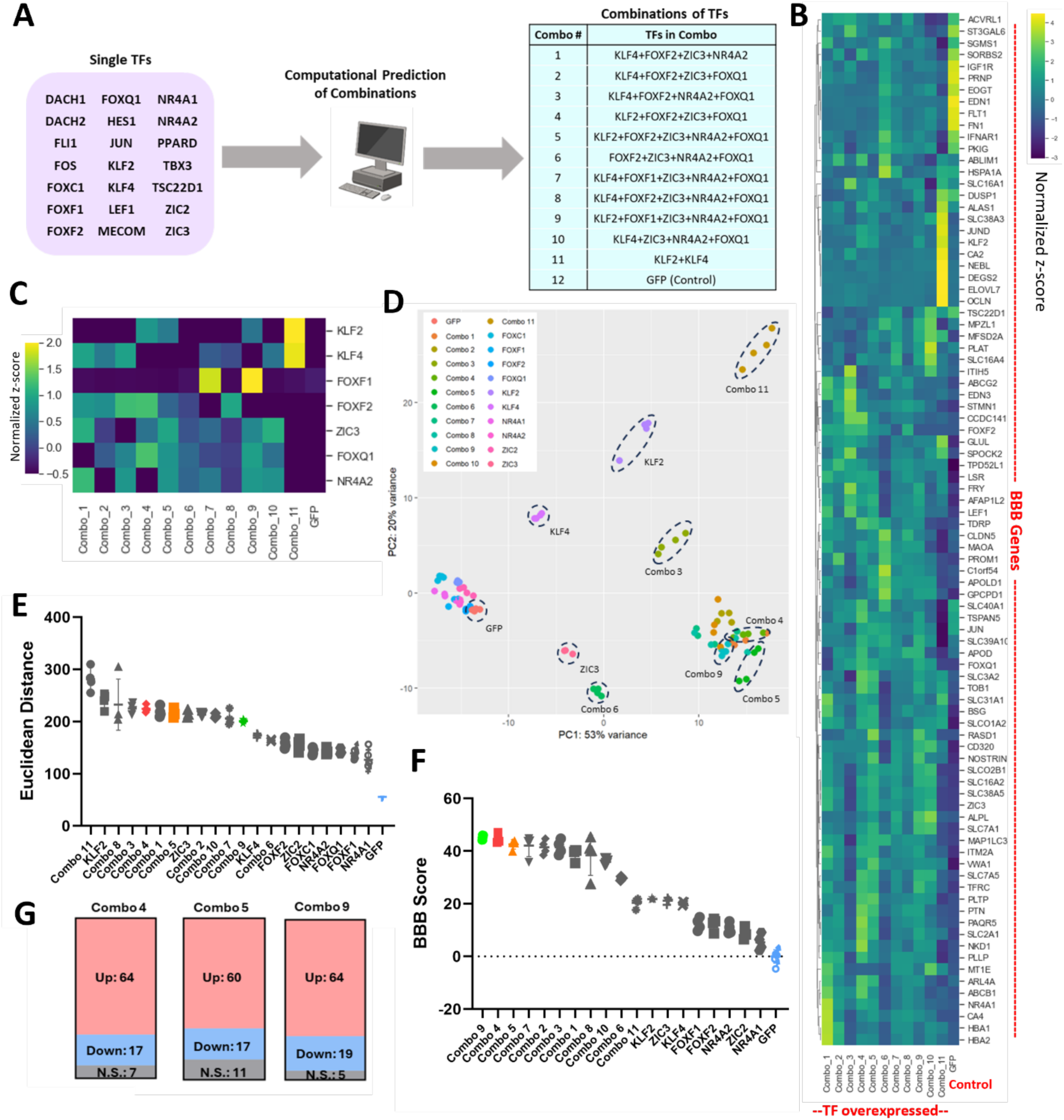
Combinatorial TF overexpression induces BBB character in fpCECs. **(A)** Schematic of the computational method to predict top TF combinations, leading to the design of 11 candidate combinations for experimental evaluation. See Methods for prediction algorithm. **(B)** Heatmap of the expression of 88 BBB-enriched genes in fpCECs overexpressing each of the 11 TF combos at MOI=2 for each TF or GFP control. Average z-score normalized to the mean expression for each gene for N=4 biological replicates is plotted. **(C)** Heatmap of the expression of 7 TFs used in combinatorial experiments. Average z-score of N=4 biological replicates is plotted. **(D)** Principal component analysis (PCA) of transcriptomes of fpCECs receiving combo TFs, single TFs, or GFP as a control. Each sample on the graphs represents an independently sequenced biological replicate. **(E)** Euclidean distances between the transcriptomes of fpCECs overexpressing the TF combos, single TFs, or GFP control. The mean±S.D. of N=4 sequencing replicates is plotted. **(F)** BBB score analysis of fpCECs overexpressing the TF combos, single TFs, or GFP control. The mean±S.D. of N=4 sequencing replicates is plotted. Colors for top TF combos in plots **(E)** and **(F)** match. **(G)** Number of 88 BBB-enriched genes upregulated, downregulated, or exhibiting no significant changes upon overexpression of TF combos 4, 5, and 9 in fpCECs. Adjusted p-value < 0.05 using the Wald test was used to determine whether upregulation or downregulation was statistically significant.

When examining BBB-like gene expression induced by TF combinations, we found that the top combinations (combos 4, 5, 9) induced expression of the efflux transporter *ABCB1* above the level observed for any single TF, while *ABCG2* was induced above GFP controls, albeit at lower levels than those observed for some single TFs. A variety of solute carrier (SLC) transporters, transferrin receptor, and junctional genes (*BSG, LSR*, and *CLDN5*) were also induced by these combinations (Fig. 6A, file S2, fig. S9, A-I). To further benchmark fpCECs overexpressing combinatorial TFs to *in vivo* brain ECs, we expanded our single cell transcriptome analysis and identified 65 brain EC-depleted genes, in addition to the 88 brain EC-enriched genes mentioned earlier (Details in Methods, gene list in file S4). We then compared the fold changes of these genes induced by combinatorial TF treatment to the gene expression differences observed *in vivo* when comparing brain ECs to non-brain ECs in mice (*30*). We found that in all three combos tested, expression of the BBB-enriched genes and BBB-depleted genes were predominantly regulated in the correct direction (Fig. 6B, fig. S10, A and B, File S5). Together these data suggest fpCECs expressing combinations of BBB-enriched TFs induce BBB-specification reminiscent to that observed when comparing *in vivo* brain ECs to non-brain ECs.

**Fig. 6.**
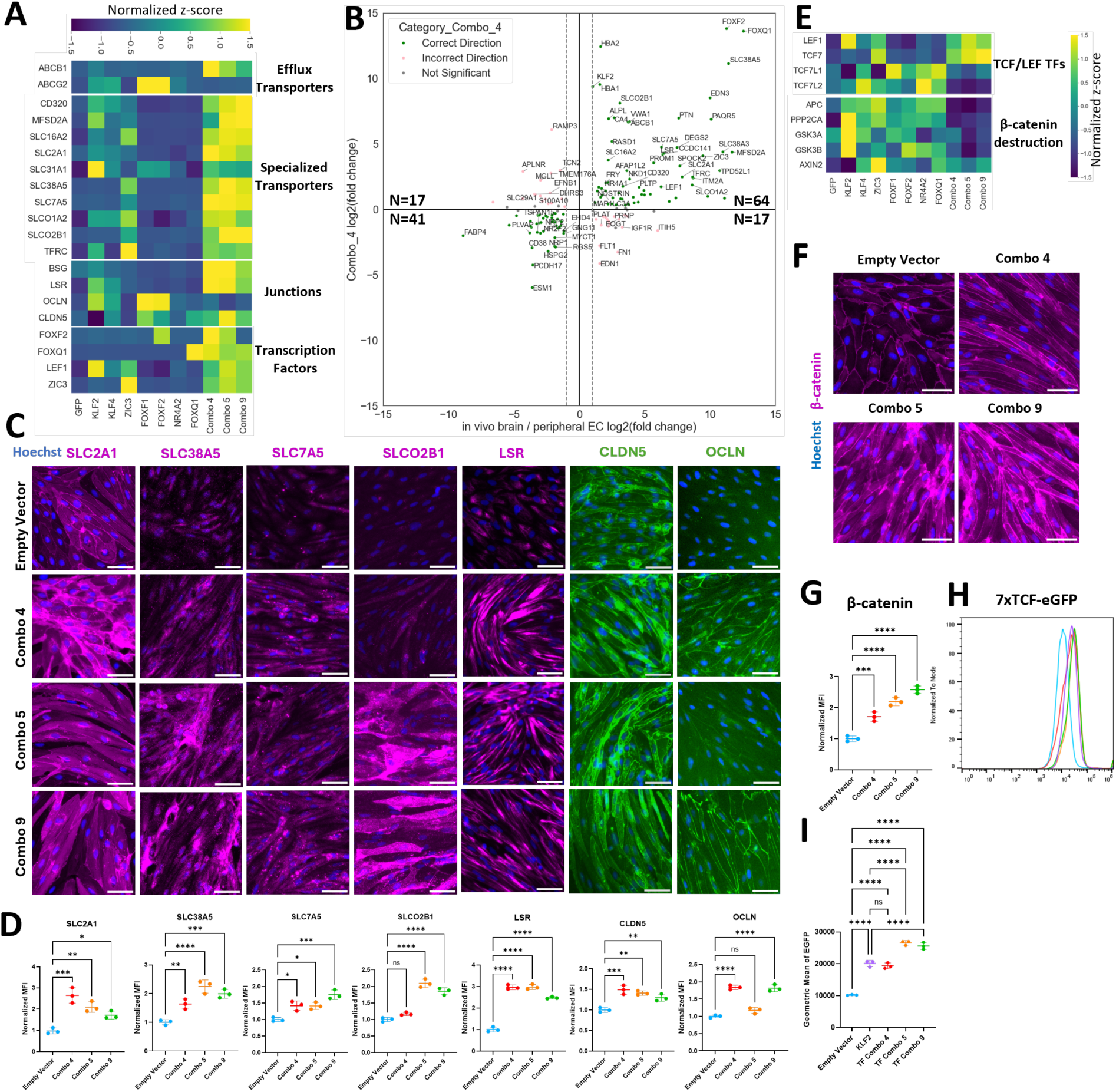
Combinatorial TF overexpression induces BBB properties in fpCECs. **(A)** Heatmap of transcript expression of efflux transporters, specialized transporters, tight junction-related genes and transcription factors that are upregulated by overexpression of TF combos 4, 5, and 9 compared to GFP control in fPCECs. Z-scores normalized to the mean expression for each gene are plotted. N=4 biological replicates per condition. **(B)** Scatter plot comparing gene expression changes elicited by Combo 4 TF overexpression to gene expression difference between brain and non-brain ECs *in vivo*. The x-axis indicates average log_2_(fold change) of gene expression in mouse brain ECs compared to liver, heart, kidney and skeletal muscle ECs for the 88 brain EC-enriched genes and 65 brain EC-depleted genes described in File S4 with known mouse-human homology. Homologous human gene names are shown. The Y-axis indicates log_2_(fold change) in Combo 4 TF-treated ECs compared to GFP-treated ECs. Dashed lines indicate log2(fold change) of 1 and –1 for *in vivo* gene expression changes. Green and pink dots represent genes significantly changing in the correct and incorrect direction, respectively, padj<0.05. Gray dots represent genes that are non-significant. Number of significant genes in each quadrant are shown. **(C)** Immunocytochemistry of specialized transporters (SLC2A1, SLC38A5, SLC7A5, SLCO2B1) and junctional proteins (LSR, CLDN5, OCLN) in fpCECs receiving either an empty vector lentivirus, or Combo 4, 5, or 9 lentiviruses. Scale bar is 50µm. **(D)** Quantification of immunocytochemistry images in **(C)**. N=3 biological replicates per condition. MFI: mean fluorescence intensity. *: p<0.05; **: p<0.01; ***:p<0.001, ****:p<0.0001 in one-way ANOVA followed by Dunnett’s post-hoc test. **(E)** Heatmap of the expression of canonical Wnt signaling TCF/LEF family TFs, and genes involved in the degradation of β-catenin. Samples are CECs receiving Combo 4, 5, 9 lentiviruses, single TFs in the aforementioned combos (*KLF2*, *KLF4*, *ZIC3*, *FOXF1*, *FOXF2*, *NR4A2*, *FOXQ1*), or GFP control lentivirus. Average of N=4 biological replicates per condition are plotted. **(F)** Immunocytochemistry of β-catenin in fpCECs receiving either an empty vector lentivirus, or Combo 4, 5, or 9 lentiviruses. Scale bar is 50µm. **(G)** Quantification of immunocytochemistry images in **(F)**. N=3 biological replicates per condition. MFI: mean fluorescence intensity. ***:p<0.001, ****:p<0.0001 in one-way ANOVA followed by Dunnett’s post-hoc test. **(H)** Flow cytometry of eGFP expression using 7xTCF-eGFP H9 (7TGP) Wnt reporter line derived EPCs transduced with either empty vector, *KLF2* or Combo 4, 5 or 9 lentiviruses with CHIR99021 treatment. **(I)** Quantification of geometric mean of fluorescence in (G). ****: p<0.0001 in one-way ANOVA followed by Tukey’s post-hoc HSD test.

We then verified elevated protein expression for a subset of these genes for TF combinations 4, 5 and 9 (Fig. 6, C and D). Increased expression was confirmed for transporters SLC2A1, SLC38A5, SLC7A5, and SLCO2B1, as well as junctional proteins LSR, CLDN5, and OCLN. Notably, OCLN was absent in our control but was induced in all cells transduced with TF combinations, indicating *de novo* acquisition of junctional OCLN expression. The fpCECs overexpressing TF combinations 4, 5 and 9 resulted in pure populations of CDH5+ ECs that retained junctional localization of CDH5 and PECAM1 (fig. S10, C and D), despite lowered transcript levels of these two genes (fig. S9, J and K). No reduction was observed in another endothelial cell marker *vWF* (fig. S9L). fpCECs overexpressing combo 4, 5, or 9 TFs, like those expressing individual TFs, retained an endothelial gene expression profile, with slight increases in mesenchymal and epithelial gene expression (fig. S10E, file S2). PACNet analysis of fpCECs overexpressing combo 4, 5, or 9 TFs also demonstrated an endothelial signature, with no clear enrichment of other cell fates (fig. S10F). As with *KLF2* alone (Fig. 3N), the TF combinations induced expression of Wnt activating TFs *LEF1* and *TCF7*, and downregulated expression of Wnt inhibitory TFs *TCF7L1* and *TCF7L2* (Fig. 6E, file S2). However, compared with individual TFs, we observed a more pronounced decrease in the expression of genes responsible for β-catenin destruction (*APC, PPP2CA, GSK3A* and *GSK3B*) (Fig. 6E), which is consistent with higher levels of canonical Wnt signaling. Thus, to confirm these transcriptomic observations, we determined that β-catenin protein levels were increased in fpCECs overexpressing TF combinations 4, 5, and 9 (Fig. 6, F and G). Also, canonical Wnt signaling was increased by TF combinations 4, 5, or 9 in the 7TGP hPSC-derived EPCs, with combinations 5 and 9 increasing Wnt signaling over *KLF2* overexpression alone (Fig. 6, H and I). These data suggest that the acquisition of BBB character in fpCECs expressing BBB-enriched TFs is at least partially mediated by increased canonical Wnt signaling.

Given that the TF combinations increased the expression of BBB genes in the developmental model of hPSC-derived CECs, we also explored the impacts of the TF combos 4, 5 and 9 on gene transcription in HUVECs. TF combinations increased transcript levels of *CLDN5*, *OCLN*, *ABCB1*, *ABCG2*, and *SLC2A1* in HUVECs, and we confirmed increased CLDN5, OCLN and SLC2A1 protein expression (fig. S11, A and B). We also evaluated whether the TF combinations would rescue to the loss of BBB properties in the immortalized human brain endothelial cell line hCMEC/D3. The gene expression changes induced by the TF combinations 4, 5, and 9 were much more modest with only increased CLDN5 gene and protein expression observed, although the CLDN5 did not appear to be correctly localized at cell-cell junctions (fig. S11, C and D). Taken together, these data suggest that the TF combinations are potent inducers of BBB properties, and their effects can be extended to non-brain endothelial cells, such as HUVECs.

### Forward programmed CNS-like ECs exhibit BBB-related phenotypes

After identifying TF combinations that induced BBB-like gene and protein expression profiles in fpCECs, we next examined whether these molecular attributes manifested in three BBB phenotypes: reduced vesicular trafficking, efflux transporter activity, and passive barrier formation. Genes associated with caveolar endocytic vesicle formation *CAV1*, *CAV2*, and *PLVAP*, were generally down-regulated by TF combos 4, 5, and 9, while *MFSD2A*, a gene enriched in CNS endothelial cells that has been linked to reduced caveolae formation and transcytosis was upregulated (*54*) (fig. S9, M-P). We further confirmed the reduction in protein expression of CAV1 and PLVAP by TF combos 4, 5 and 9 (Fig. 7, A and B). To capture distinct vesicular uptake mechanisms in fpCECs, we tested both albumin and 10 kDa dextran accumulation. Albumin is a serum protein trafficked across the BBB primarily via caveolin-mediated endocytosis, whereas 10 kDa dextran is internalized mainly by nonspecific fluid-phase pinocytosis and thus serves as a tracer of general endocytic uptake (*55*, *56*). TF combos 4, 5, and 9 all resulted in a significant decrease in intracellular accumulation of both albumin and 10 kDa dextran at 37°C, but no difference was observed at 4°C as this condition limits membrane fluidity and inhibits endocytosis (Fig. 7, C-F). Together, these data suggested that fpCECs demonstrated both reduced caveolae-mediated uptake and general fluid phase endocytosis.

**Fig. 7.**
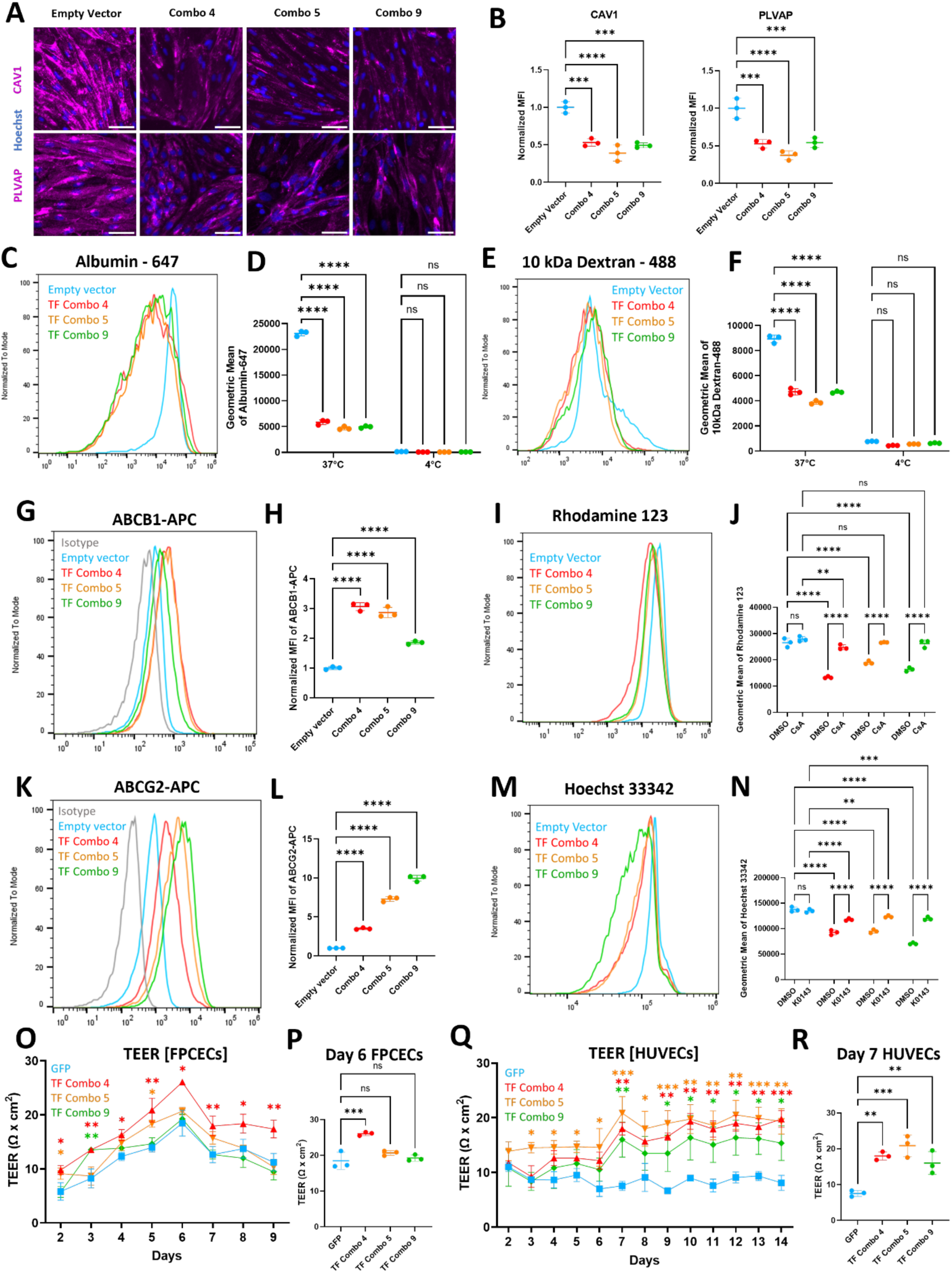
Combinatorial TF overexpression generates fpCECs that exhibit key BBB phenotypes. **(A)** Immunocytochemistry of CAV1 and PLVAP in fpCECs receiving either an empty vector lentivirus, or TF combos 4, 5, or 9 lentiviruses. Scale bar is 50µm. **(B)** Quantification of CAV1 and PLVAP immunocytochemistry images. N=3 biological replicates per condition. MFI: mean fluorescence intensity. ***:p<0.001; ****:p<0.0001 in one-way ANOVA followed by Dunnett’s post-hoc test. **(C)** Flow cytometry analysis of endocytosed albumin – Alexa Fluor 647 in fpCECs receiving either TF combos 4, 5, or 9 lentiviruses, or an empty vector lentivirus control. Representative histograms are from the 37 °C condition. **(D)** Quantification of geometric mean of albumin – Alexa Fluor 647 from flow cytometry analysis of fpCEC populations receiving either TF combo 4, 5, or 9 lentiviruses, or an empty vector lentivirus control. Experiments were performed at 37 °C or 4 °C. N=3 biological replicates for each condition. Legends in (C) apply to (D). ****:p<0.0001 in two-way ANOVA followed by Tukey’s post-hoc test. **(E)** Flow cytometry analysis of endocytosed 10kDa dextran – Alexa Fluor 488 in fpCECs receiving either TF combos 4, 5, or 9 lentiviruses, or an empty vector lentivirus control. Representative histograms are from the 37 °C condition. **(F)** Quantification of geometric mean of 10kDa dextran – Alexa Fluor 488 from flow cytometry analysis of fpCEC populations receiving either TF combo 4, 5 or 9 lentiviruses, or an empty vector lentivirus control. Experiments were performed at 37 °C or 4 °C. Legends in (E) apply to (F). N=3 biological replicates for each condition. ****: p<0.0001 in two-way ANOVA followed by Tukey’s post-hoc HSD test. **(G)** Flow cytometry analysis of ABCB1 expression of fpCECs receiving either TF combo 4, 5, or 9 lentiviruses, or an empty lentivirus control. **(H)** Quantification of geometric mean of ABCB1 expression from flow cytometry analysis of fpCEC populations receiving either TF combo 4, 5 or 9 lentiviruses, or an empty lentivirus control. ****: p<0.0001 in one-way ANOVA followed by Dunnett’s post-hoc test. **(I)** Flow cytometry analysis of Rhodamine 123 (Rh123), a substrate of ABCB1. Flow histograms represent fpCECs receiving either TF combos 4, 5, or 9 lentiviruses, or an empty lentivirus control. **(J)** Quantification of geometric mean of Rhodamine 123 from flow cytometry analysis of fpCEC populations receiving either TF combo 4, 5 or 9 lentiviruses, or an empty vector lentivirus control, in the presence or absence of the ABCB1 inhibitor CsA. Mean ± SD, **:p<0.01, ****: p<0.0001 in two-way ANOVA followed by Tukey’s post-hoc HSD test. **(K)** Flow cytometry analysis of ABCG2 expression of fpCECs receiving either TF combos 4, 5, or 9 lentiviruses, or an empty vector lentivirus control. **(L)** Quantification of geometric mean of ABCG2 expression from flow cytometry analysis of fpCEC populations receiving either TF combos 4, 5 or 9 lentiviruses, or an empty vector lentivirus control. ****: p<0.0001 in one-way ANOVA followed by Dunnett’s post-hoc test. **(M)** Flow cytometry analysis of Hoechst 33342, a substrate of the BCRP efflux pump. Flow histograms represent fpCECs receiving either TF combos 4, 5, or 9 lentiviruses, or an empty vector lentivirus control. **(N)** Quantification of geometric mean of Hoechst 33342 from flow cytometry analysis of fpCEC populations receiving either TF combos 4, 5, or 9 lentiviruses, or an empty vector lentivirus control, in the presence or absence of Ko143. Mean ± SD are shown, **:p<0.01, ***:p<0.001, ****: p<0.0001 in two-way ANOVA followed by Tukey’s post-hoc HSD test. **(O)** Transendothelial electrical resistance (TEER) of fpCECs receiving either TF combos 4, 5, or 9 lentiviruses, or control GFP lentivirus. Cells were seeded on Day 0 and transduced on Day 1. TEER measurement started on Day 2, 24 hours after transduction, Mean ± SD of N=3 biological replicates were plotted. *: p<0.05; **: p<0.01 in one-way ANOVA followed by Dunnett’s post-hoc test. **(P)** Quantification of TEER measurement on day 6 of fpCECs receiving either TF combos 4, 5, or 9 lentiviruses, or GFP control lentivirus. Mean ± SD of N=3 biological replicates were quantified, **: p<0.01; ***: p<0.001 in one-way ANOVA followed by Dunnett’s post-hoc test. **(Q)** Transendothelial electrical resistance (TEER) of HUVECs receiving either TF combo 4, 5, or 9 lentiviruses, or GFP lentivirus. TEER measurement started on Day 2, 24 hours after transduction, Mean ± SD of N=3 biological replicates were plotted. *: p<0.05; **: p<0.01; ***: p<0.001 in one-way ANOVA followed by Dunnett’s post-hoc test. **(R)** Quantification of TEER measurement on day 7 of HUVECs receiving either TF combos 4, 5, or 9 lentiviruses, or GFP control lentivirus. Mean ± SD of N=3 biological replicates were quantified **:p<0.01, ***: p<0.001 in one-way ANOVA followed by Dunnett’s post-hoc test.

Another key feature of BBB ECs is efflux transporter activity with ABCB1 (p-glycoprotein) and ABCG2 (BCRP) playing key roles. Concordant with transcriptional induction of *ABCB1* by TF combinations (fig. S9Q), flow cytometry indicated significant expression of cell surface ABCB1 protein expression induced by the TF combinations (Fig. 7, G and H). To evaluate whether the increase in expression corresponded to efflux function, we measured the accumulation of Rhodamine123 (Rh123), a substrate of p-glycoprotein (P-gp), in fpCECs with or without P-gp inhibitor cyclosporin A (CsA). We observed that fpCECs transduced with combos 4, 5 and 9 had reduced Rh123 cellular accumulation compared to control CECs (Fig. 7, I and J), indicating that these cells actively effluxed Rh123. By contrast, when fpCECs were treated with CsA, the decrease in Rhodamine123 accumulation was abrogated to the level of the control CECs, whereas control CECs possessed no observable ABCB1 efflux activity (Fig. 7J). The fpCECs transduced with combos 4, 5, and 9 also possessed an increased cell surface ABCG2 protein expression that correlated with transcript levels (Fig. 7, K and L, fig. S9R). To verify ABCG2 functionality, Hoechst 33342 was used as the ABCG2 substrate, and a reduction in Hoechst accumulation was observed in fpCECs transduced with combos 4, 5, and 9 compared to control CECs, indicating these cells actively effluxed Hoechst 33342 (Fig. 7M). When fpCECs were treated with ABCG2 inhibitor Ko143, increased Hoechst 33342 accumulation was observed, whereas control CECs possessed no observable ABCG2 efflux activity (Fig. 7N). Together, these data indicate that while control CECs possessed no detectable efflux activity, fpCECs transduced with combos 4, 5, and 9 exhibited robust ABCB1 and BCRP activity.

The third key BBB property measured was tight junction barrier formation using transendothelial electrical resistance (TEER). We observed that the fpCECs transduced with TF combination 4 had a modest increase in TEER over the CEC control (Fig. 7, O and P). TF combinations 5 and 9 did not show increased TEER despite also having increased CLDN5 protein expression and observable OCLN expression (Fig. 6C). Since lentiviral transduction itself had a deleterious impact on TEER in hPSC-derived CECs (fig. S12A), we tested TEER response to lentivirally administered TF combinations in HUVECs where lentivirus did not impact TEER (fig. S12, A and B). TF combinations 4, 5, and 9 all increased TEER in HUVECs (Fig. 7, Q and R). These increases correlated with the increased CLDN5 protein expression along with the appearance of OCLN protein expression in HUVECs (fig. S10, A and B).

## Discussion

In this study, we adopted a comprehensive transcriptomics-based approach to identify and validate BBB-enriched transcriptional regulators that drive brain endothelial genotype and phenotype acquisition in hPSC-derived CECs as a human developmental model of the BBB. Specifically, we discovered that overexpression of transcription factors *KLF2*, *KLF4*, *FOXF1*, *FOXF2*, *ZIC2*, *ZIC3*, *NR4A1*, *NR4A2*, *FOXC1* or *FOXQ1* induced subsets of BBB-enriched transcripts. Combinations of these TFs led to synergistic improvements in BBB-specific character in terms of transcript and protein expression as well as BBB function. Taken together, the identified TFs and their combinations orchestrate BBB specification in hPSC-derived CECs.

Among the TFs, *KLF2* and *KLF4* are potent inducers of BBB-like gene expression in hPSC-derived CECs. Additionally, *KLF2* and *KLF4* induced expression of other BBB-enriched TFs, such as *FOXF2*, *FOXQ1*, *LEF1* and *ZIC3*, suggesting that they might be upstream regulators of BBB specification in developing ECs. The effects of *KLF2* and *KLF4* overexpression on BBB gene transcription worked in synergy with Wnt activation. Analysis of Wnt signaling components at the transcriptional level revealed that the synergy is accompanied by a switch from inhibitory *TCF7L1/TCF7L2* expression to *LEF1/TCF7* activation, leading to a further induction of canonical Wnt signaling in a Wnt reporter line. Given the importance of Wnt signaling in BBB development and maintenance (*11*), the ability of *KLF* TFs to enhance Wnt activation may further drive BBB specification. These findings are especially intriguing due to the role of *KLF2* and *KLF4* in vascular physiology. In endothelial cells, shear stress induces *KLF2* and *KLF4* expression (*48*). Shear stress is established when nascent capillaries become perfused and is considered to be a crucial factor for endothelial physiology commonly absent in static *in vitro* cell culture (*57*). However, shear stress cannot be the sole contributor to BBB-specific gene expression because shear stress is ubiquitous in the vasculature throughout the body, yet the tight endothelial barrier phenotype is only present at the BBB or blood-retina barrier. Moreover, *KLF2* and *KLF4* are enriched at the BBB compared with peripheral vasculature, suggesting that they could play unique roles in BBB specification in addition to more general shear response mechanisms (Fig. 1). *Klf2* deletion is embryonic lethal (*58–61*), and double endothelial-specific knockout of *Klf2* and *Klf4* in adult mice causes widespread vascular dysfunction including myocardial infarction and cerebral hemorrhage, highlighting a key role for KLF family TFs in vascular function (*62*). In addition, *KLF2*-deficient mice have been found to have increased infarct volume following a stroke and increased BBB permeability (*63*), suggesting that *KLF2* can specifically impact BBB properties *in vivo*. Future studies are needed to further understand the impact of *KLFs* on BBB development and maintenance.

We also analyzed the detailed global transcriptional effects elicited by a broad set of BBB-enriched TFs, with a focus on *FOXF1*, *FOXF2*, *ZIC2*, *ZIC3*, *FOXQ1*, *FOXC1*, *NR4A1*, and *NR4A2*. Although the overall effects of these TFs on the expression of BBB-enriched genes were not as significant as *KLF2* or *KLF4*, they induced discrete subsets of BBB genes. Notably, *FOXF1* and *FOXF2* induced expression of several specialized transporters such as *SLC2A1, SLC39A10, SLC41A1*, and genes associated with lipid metabolism at the BBB including *ELOVL7, GPCPD1* and *PLTP*. *Foxf2* has been shown to play a role in BBB formation and maintenance, as knockout mice have BBB dysfunction, BMEC thickening and increased transcytosis (*64*). Recently, it has been found that *Foxf2* is involved in the maintenance of brain endothelial cell function in adult mice (*65*). In addition, while *FOXQ1*, *ZIC3* and *NR4A2* have been shown to be enriched in BMECs (*29*, *66*), their function in controlling BBB properties within BMECs has not yet been explored in detail. In this study, *FOXQ1* and *FOXC1* reduced caveolin-1 expression, *ZIC2* and *ZIC3* significantly induced the expression of several BBB-enriched SLC transporters and *NR4A1* and *NR4A2* modestly induced expression of several BBB-specific transporters. Corroborating our findings that these TFs can drive acquisition of BBB properties *in vitro*, overexpression of *FOXF2* and *ZIC3* in hPSC-derived ECs led to enhanced tight junction protein expression (*67*), and overexpression of *FOXF2* and *ZIC3* in HUVECs induced expression of some BBB genes (*29*). The rich data sets provided in this study will support future exploration of the impact of these TFs in the acquisition and maintenance of BBB properties both *in vitro* and *in vivo*.

Given that overexpression of individual TFs led to induction of different subsets of BBB genes, we tested combinatorial overexpression of TFs to capture a more comprehensive BBB-like gene expression profile. Specifically, combinations of *KLF2* or *KLF4* with *FOXF2*, *ZIC3*, *NR4A2*, and *FOXQ1* enhanced the expression of 62 out of 88 BBB-enriched genes, leading to a higher BBB score than fpCECs overexpressing individual TFs. We also found that overexpressing combinatorial TFs in fpCECs elicited changes in gene expression that mimic the differences observed between brain and peripheral ECs *in vivo.* These data suggest that TF combinations provide many of the unique cues that distinguish brain ECs from peripheral ECs *in vivo*. Functionally, we demonstrated that fpCECs overexpressing combinatorial TFs possess key BBB properties such as greatly reduced endocytosis, P-gp and BCRP efflux transport activity, and improvements in tight junction protein expression leading to modest improvements in barrier integrity. Additionally, we evaluated the effect of overexpression of the top 3 TF combinations in 2 other endothelial cells lines, non-barrier forming primary HUVECs, and the hCMEC/D3 immortalized brain endothelial cell line. The forward programmed HUVECs showed increased tight junction protein expression of CLDN5 and OCLN, along with increased TEER values, showcasing the potential of these TF combinations to induce BBB properties in non-barrier forming endothelial cells. The hCMEC/D3 cell line has a limited barrier phenotype as a result of immortalization and extended *in vitro* culture, with low expression of the tight junction proteins CLDN5 and OCLN (*68*). Overexpressing the TF combinations in hCMEC/D3 cells increased CLDN5 expression, which shows the ability of the TFs to partially rescue the loss of tight junction phenotype. Together, these data demonstrate that the effect of TF combos can be extended beyond hPSC-derived endothelial systems, further validating that these TFs can induce BBB phenotypes in cells that lack these properties.

Finally, we envision that fpCECs transduced with TF combinations 4, 5, or 9 could serve as *in vitro* BBB models for a variety of applications. While the induction of additional BBB properties by TF overexpression in hPSC-derived CECs is tantalizing, there are several technical hurdles remaining. In this study, we used lentivirus to deliver TFs under a constitutive promoter. This approach allowed for rapid screening to identify candidate TFs but does not allow for precise control of dose and timing of TF overexpression. Moreover, passive barrier properties were negatively impacted by lentiviral transduction, so alternative gene delivery strategies may be needed to maximize the TF impact on certain BBB properties. For instance, we envision that utilizing inducible promoters in hPSC lines with stably integrated genetic constructs will allow tuning of the resulting fpCECs. Although the induction of BBB character was substantial, it is important to note that not all BBB properties were induced to the appropriate extent by the tested TF combinations. Thus, other factors, such as additional TFs or interactions with other neurovascular unit cell types, will be necessary to further optimize the fpCEC product.

## Materials and Methods

### Maintenance of hPSCs

Human pluripotent stem cells (IMR90-4 from WiCell WISCi004-B and H9 7TGP Wnt reporter line) were maintained on Matrigel (Corning 356234) coated plates in TeSR-E8 media (STEMCELL Technologies). hPSCs were passaged using Versene (Gibco 15040066) at 1:6 ratio upon reaching 70% confluency. Regular mycoplasma and G-band karyotyping tests were performed.

### Differentiation of Forward-programmed CECs

Differentiation was performed according to a slightly modified version of our previously published methods (*25*, *42*). Briefly, on day –3, singularized IMR90-4 hPSCs were seeded onto 12-well plates (Corning 3513) coated with Matrigel at a density of 30,000 cells per well in E8 media. On day –2 and –1, medium changes with mTeSR were performed. On day 0 and day 1, cells were treated with 6 μM CHIR99021 (Tocris 4423) for 48 hours in LaSR medium prepared according to our previous protocol (*27*). From day 2 to day 5, cells were treated with LaSR medium with 50 ng/mL VEGF (Peprotech 100-20). On day 5, typically 15% to 30% of cell populations are CD31+ EPCs. On day 5, EPCs were purified by magnetic sorting of CD31+ cells, using anti-CD31-biotin antibody (Miltenyi 130-110-667), anti-biotin microbeads (Miltenyi 130-097-046) and a QuadroMACS separator (Miltenyi 130-091-051). Magnetic sorting was performed according to manufacturer’s recommendations.

Purified CD31+ EPCs were further differentiated to CECs according to our previously published method (*27*). Briefly, Collagen IV (Sigma-Aldrich C5533) was used to coat tissue-culture plates at 10 ug/cm^2^. hECSR media was prepared by mixing hESFM (Gibco 11111044) with 2% B-27 (Gibco 17504044) and 20ng/mL FGF2 (Peprotech 100-18B). EPCs obtained as described above were suspended in hECSR medium supplemented with 4-6 µM CHIR99021 (Tocris 4423) and plated at approximately 3 × 10^4^ cells/cm^2^. Lentiviruses delivering TFs, GFP control, or empty vector control were dosed the following day (day 6) at MOI=2 for each lentivirus. For transduction of TF combinations, each of the TFs which are part of the combination were dosed at MOI=2. For assays with fluorescence readouts overlapping with the GFP channel, the empty vector control lentivirus was used in place of GFP lentivirus. Media was changed every 48 hours. fpCECs were analyzed after 5 days in culture for all assays, except for TEER, wherein the cells were cultured beyond this timepoint.

### TF plasmids and lentivirus production

TF lentivirus transfer plasmids were obtained from Addgene (Table S2) as part of the MORF library lentivirus collection (*19*) as gifts from Feng Zhang. To generate the empty vector control, the GFP coding sequence was excised from the GFP plasmid. The lentiviral packaging plasmids psPAX2 (Addgene 12260) and pMD2.G (Addgene 12259) were obtained from Addgene as gifts from Didier Trono. To produce lentivirus, Lenti-X 293T cells (Takara 632180) were maintained on collagen I-coated (Gibco A1048301) six-well plates in DMEM (Life Technologies 11965092) supplemented with 10% FBS (Peak Serum PS-FB1), 1 mM sodium pyruvate (Life Technologies 11360070), and 0.5× GlutaMAX Supplement (Life Technologies 35050061). When 293T cells reached 90% confluence, psPAX2 (1 μg per well), pMD2.G (0.5 μg per well), and lentivirus transfer plasmids (1.5 μg per well) were co-transfected using FuGENE HD Transfection Reagent (9 μl per well) (Promega E2311). Medium was replaced 16 hours after transfection, and virus-containing supernatants were collected 24, 48, and 72 hours later. Supernatants were filtered through a 0.45-μm filter and concentrated 100× using Lenti-X Concentrator (Takara 631231). Titer of lentivirus was determined using Lenti-X GoStix Plus (Takara 631280) according to manufacturer’s recommendations.

### Culture conditions for HUVECs and hCMEC/D3

The human umbilical vein endothelial cells (HUVECs) and human cerebral microvascular endothelial cells/clone D3 (hCMEC/D3) were grown in EBM-2 medium (Lonza, Switzerland) supplemented with EGM-2 Endothelial SingleQuots Kit (Lonza). Cells were grown on 6-well plates (Corning) coated with rat tail collagen type-I. Cells were passaged using Accutase (STEMCELL Technologies) at a 1:6 ratio upon reaching 80-90% confluence. Cells were maintained in the complete EGM2 media (EBM2 basal media + supplements) with medium replacement every other day. Cells were plated at a density of approximately 6 × 10^4^ cells/cm^2^ and 3 × 10^5^ cells/cm^2^ for qPCR and TEER assays respectively. Cells were transduced the next day with lentivirus carrying the GFP control or TFs at MOI=2 and analyzed after 5 days in culture, except for TEER experiments, wherein the cells were cultured for longer.

### Single cell RNA sequencing analysis

We obtained single cell RNA-seq gene expression matrices from GEO or ArrayExpress (GSE134355, GSE76381, GSE119212, GSE106118, GSE124395, GSE130646, GSE175895, E-MTAB-8077). We used R (version 3.6.2) and the Seurat package (*69–73*) for all analyses. Data was normalized to 2000 highly variable features (genes) for each dataset independently. We performed principal component analysis and used the first 30–50 principal components to perform UMAP embedding. For marker identification, we used the FindAllMarkers function, and selected genes with adjusted P-values < 0.05 based on the default Wilcoxon rank sum test (a non-parametric test that does not assume normally-distributed data) with Bonferroni correction. We used the Seurat functions DimPlot, FeaturePlot, VlnPlot, DoHeatmap, and DotPlot for visualization. To integrate different human brain scRNA-seq datasets, we used methods previously developed (*40*). We used FindMarkers function from Seurat to perform differential expression tests, and crossed the list of differentially expressed genes with a published list of human TFs to identify target TFs(*74*). Statistical analyses of differential expression used the Wilcoxon rank sum test with Bonferroni correction.

### Bulk RNA sequencing library preparation and bioinformatics analysis

RNA-seq was performed on fpCECs derived from the IMR90-4 hPSC line. Four independent biological replicates were generated as described in previous section. Cells were resuspended in Tri Reagent (Zymo R2050-1-200) for cell lysis. RNA was then extracted using Direct-zol RNA Miniprep kit (Zymo R2050) according to manufacturer’s guidelines. RNA quality was assayed using Agilent 2100 Bioanalyzer with Agilent RNA 6000 Pico Kit (Agilent 5067-1513). All samples have RIN number larger than 9. Poly A-enriched mRNA from 1000ng of total RNA was then prepped using the NEBNext Poly(A) mRNA Magnetic Isolation Module (NEB E7490L) on a DynaMag-96 Side Magnet (Invitrogen 12331D). 1st strand cDNA synthesis and sequencing library preparation was performed using xGen RNA Library Preparation Kit (IDT 10009814), with library purification steps done with AMPure XP beads (Beckman Coulter A63882). Quality of the sequencing library was checked on an Agilent 2100 bioanalyzer using High Sensitivity DNA Kit (Agilent 5067-4626). DNA concentration of sequencing library was quantified on Qubit 3 with dsDNA high sensitivity kit (Invitrogen Q32851). Sequencing was performed on a NovaSeq X (Illumina), with approximately 30 million 150 bp paired end reads obtained for each sample.

FASTQ files were first tail-trimmed using Trimmomatic (*75*). The trimmed reads were then aligned to the human genome (hg38) using RNA STAR (*76*). Gene counts were summarized using featureCounts (*77*). Gene counts were fed into DESeq2 (*78*) implemented in R for differential expression analysis. The Wald test with Benjamini–Hochberg correction was used to generate adjusted p-values. Principal component analysis was performed on counts after the DESeq2 variance stabilizing transformation. Heatmap was generated based on TPM values using seaborn (*79*) in Python.

### Computational processing of transcriptomics data

To quantify the level of changes induced by overexpression of each TF, we calculate the Euclidean distance between the TF-overexpression transcriptome profile and the average of GFP transcriptome profile. Euclidean distance is calculated as the sum of squares of differences of the natural log of TPM values for each gene.

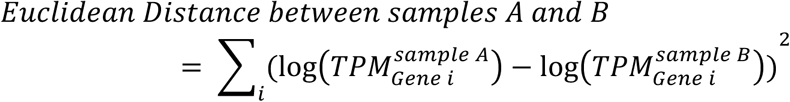

To quantify overall BBB gene expression, we created a custom variable: the BBB score. It is defined as the sum of log_2_ (fold changes) of the 88 BBB enriched genes (Fig. 2D). The list of 88 genes were curated from single cell RNA-seq datasets, for genes that were significantly upregulated more than two-fold in murine brain ECs compared to peripheral ECs sequenced by Kalucka et al., (*30*), while still having significant expression levels (TPM>10) in human brain ECs sequenced by Crouch et al., (*41*). Since overexpression of TFs can increase or decrease the expression of these 88 genes, BBB score captures both the benefits and disadvantages. However, to limit the influence of large fold change of a single gene on BBB score, we capped the contribution from each of the 88 genes from –1 to 1. In summary, the BBB score is calculated as:

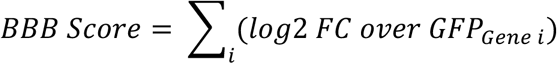

To predict combinations of TFs that capture a more *in vivo* BBB-like phenotype, we first calculated the change induced by each of the TFs, and called the variable Δ *TF_i_*. Then we used the pseudobulk function from the glmGamPoi package(*80*), and obtained gene counts from the three single cell RNA-seq analyses done in Fig. 1. We then calculated the transcriptomic difference between the BBB-like EC population and the non-BBB EC population, and called the variable Δ *in vivo*. We then calculated the average Δ *TF_i_* profiles composed of four or five TFs, and calculated the Pearson distance from these average Δ *TF_i_* profiles to Δ *in vivo* using the dist function from the Python math package (*81*), mirroring a previously published approach (*19*). We then ranked the distances and tabulated the top 20 predictions with smallest Pearson distances between Δ *in vivo* and averaged Δ *TF_i_* profiles. The number of TF appearances in these top 20 four-TF combinations and top 20 five-TF combinations were counted. Since top predicted combinations are predominantly permutations containing ZIC family TFs, KLF family TFs, FOXF family TFs, *FOXQ1* and *NR4A2*, we then designed 11 TF combinations that included these top TFs for experimental testing (Fig. 5A).

To quantitatively analyze the effects of combinatorial TF treatment on fpCECs, we also mined mouse single cell RNA-seq datasets for genes depleted in brain ECs, compared to peripheral ECs in the liver, kidney, heart and skeletal muscle (*30*). The analysis identified 65 genes significantly downregulated more than two-fold in murine brain ECs compared to peripheral ECs in the Kalucka et al. dataset (*30*), while still retaining expression (TPM>1) in human brain ECs in the Crouch et al dataset (*41*). Using the Kalucka et al. dataset, we then pseudobulked the sequenced cells using Seurat, segregrating with regard to conditions (brain and non-brain) to generate gene counts (*69*). Fold changes induced by combinatorial TF treatment were plotted against *in vivo* differences between mouse brain ECs and non-brain ECs using Seaborn (*79*).

### Immunocytochemistry

Cells were fixed with –20°C methanol (Sigma, 67-56-1) or 4% paraformaldehyde (PFA; Electron Microscopy Sciences, 15700) in DPBS (Gibco 14190144) for 15 min at room temperature. Following three washes in DPBS the cells were blocked in 10% goat serum (Sigma, G9023) for 30 min. Cells were then incubated with primary antibodies at indicated dilution ratios at 4 °C overnight (Table S3). Cells were washed with PBS three times followed by incubation with secondary antibodies (Table S3) and 20uM Hoechst 33342 (Thermo Scientific, 62249) for 1 hour at room temperature. Cells were then washed three times. For widefield fluorescence microscopy, we used an Eclipse Ti2-E epifluorescence microscope (Nikon, Japan). For quantification of immunocytochemistry images, background signals from a secondary antibody only control were deducted from all conditions.

### RT-qPCR

RNA was extracted using Direct-zol RNA Miniprep kit (Zymo, R2050). Reverse transcription was performed to obtain cDNA using GoScript Reverse Transcriptase with Oligo(dT) kit (Promega, A2791). Real-time gene expression analysis was conducted using 20-μl reactions containing Luna Universal qPCR Master Mix (NEB, M3003) along with primers specific for genes of interest (Table S4). PCR was run according to manufacturer protocols on Agilent AriaMx Real-Time PCR system. Gene expression (2^−ΔΔCt^) was normalized to *HPRT1*.

### Flow cytometry

Cells were singularized with Accutase for 7 min at 37°C. Cell suspensions were passed through 40 µm cell strainers into 4× volume of DMEM/F-12 (Gibco 11320033) and centrifuged for 5 min, 200× g. Live cells were resuspended in flow buffer, composed of DPBS (Gibco 14190144) with 0.5% bovine serum albumin (Sigma A2153) and 2 mM EDTA (Sigma; 03620) containing pre-conjugated antibody and incubated at 4°C for 15 min. Cells were then pelleted and washed three times with flow buffer. Samples were analyzed using the Attune NxT V6 flow cytometer (ThermoFisher) and results were analyzed using the FlowJo software.

### Dextran and albumin endocytosis assays

We used two different fluorescently labeled tracers previously reported to be associated with caveolae-mediated and fluid-phase endocytosis: Alexa Fluor 647-conjugated albumin (Invitrogen A34785) and Alexa Fluor 488-conjugated 10kDa dextran (Invitrogen D22910). Dextran was added at 10 µM to the medium of day 5 fpCECs. Plates were incubated on rotating platforms at 37 or 4°C for 2 hr. Medium was removed and cells were washed once with DPBS, and then incubated with Accutase for 7 min at 37°C. Cell suspensions were passed through 40 µm cell strainers into 4× volume of DMEM/F-12 and centrifuged for 5 min, 200× g. Pellets were resuspended in flow buffer and analyzed on a Attune flow NxT V6 flow cytometer (ThermoFisher). FlowJo software was used to analyze flow cytometry results.

### P-gp and BCRP efflux transport activity assays

We used Rhodamine123 (Rh123) as a substrate of P-gp, and cyclosporine-A (CsA) as an inhibitor of P-gp. Day 5 fpCECs were incubated with 10 µM Rh123 for 2 h at 37°C, followed by 1 h incubation in media containing 10µM Rh123+ 10 µM CsA or Rh123 + DMSO. Cells were washed once with DPBS and incubated with Accutase for 7 minutes at 37°C. Cell suspensions were passed through 40 µm cell strainers into 4× volume of DMEM/F-12 and centrifuged for 5 min, 200× g. Pellets were resuspended in flow buffer and Rh123 intensity was analyzed on a Attune NxT V6 flow cytometer (ThermoFisher). FlowJo software was used to analyze flow cytometry results.

For the BCRP assay, we used Hoechst 33342 as a substrate of BCRP and Ko143 as an inhibitor. Day 5 fpCECs were incubated with 10nM Hoechst for 2 h at 37°C, followed by 1 h incubation in media containing 10nM Hoechst + 10 µM Ko143 or 10nM Hoechst + DMSO. Cells were washed once with DPBS and processed as above to measure Hoechst intensity using the Attune NxT V6 flow cytometer.

### Transendothelial electrical resistance (TEER)

Transwell inserts (6.5 mm diameter with 0.4 µm pore polyester filters) (Corning) were coated with 50 µL of a solution of collagen IV (400 µg/mL) and fibronectin (100 µg/mL) in water for 4 h at 37°C. EPCs were seeded (Day 0) on Transwell inserts at 10^5^ cells/cm^2^ in hECSR medium supplemented with 4-6 µM CHIR. 200 µL media was added to the apical chamber and 800 µL to the basolateral chamber. Cells were transduced the next day (Day 1), and TEER was measured daily starting Day 2 using an EVOM2 epithelial voltohmmeter with STX2 chopstick electrodes (World Precision Instruments, Sarasota, FL). Medium was replaced every other day. TEER values were corrected by subtracting the resistance of a blank collagen IV/fibronectin-coated Transwell insert without cells and multiplying by the filter surface area of 0.33 cm^2^.

### Statistical Analysis

One-way analysis of variance (ANOVA) was used for comparison of means from three or more experimental groups. Two-way ANOVA was used for comparison of means where there were two independent variables influencing the outcome. Following ANOVA, Dunnett’s post hoc test was used for comparison of multiple treatments to a single control, or Tukey’s Honestly Significant Difference (HSD) test was used for multiple pairwise comparisons. For scRNA-seq datasets, Wilcoxon rank sum test was used for differential gene expression analysis. For bulk RNA-seq datasets, Wald test was used for differential gene expression analysis. Details of replicates and significance are provided in figure legends.

## Author contributions

Conceptualization: S.T., Y.D., S.P.P., E.V.S.

Methodology: S.T., Y.D., F.Y.H., S.M.B.

Investigation: S.T., Y.D., F.Y.H., S.M.B, R.D., S.P.P., E.V.S

Visualization: S.T., Y.D.

Supervision: S.P.P., E.V.S

Writing—original draft: S.T., Y.D.

Writing—review & editing: S.T., Y.D., S.P.P., E.V.S

S.T. and Y.D. designed, performed, and analyzed experiments. S.T. and Y.D. wrote the manuscript. S.T., Y.D., and S.M.B. analyzed scRNA-seq datasets. F.Y.H. performed experiments. E.V.S. S.P.P, and R.D. designed and supervised the overall study. All authors read and approved the final manuscript.

## Data and materials availability

RNA-seq data has been deposited in the Gene Expression Omnibus (GEO). All data needed to support the conclusions in this paper is available in the main text or the supplementary materials.

## Supplementary Materials

See Supplementary Materials document.

## Supplementary Materials for

### This PDF file includes

Figs. S1 to S12

Tables S1 to S4

Legends for Files S1 to S5

### Other Supplementary Materials for this manuscript include the following

Files S1 to S5

**Fig. S1.**
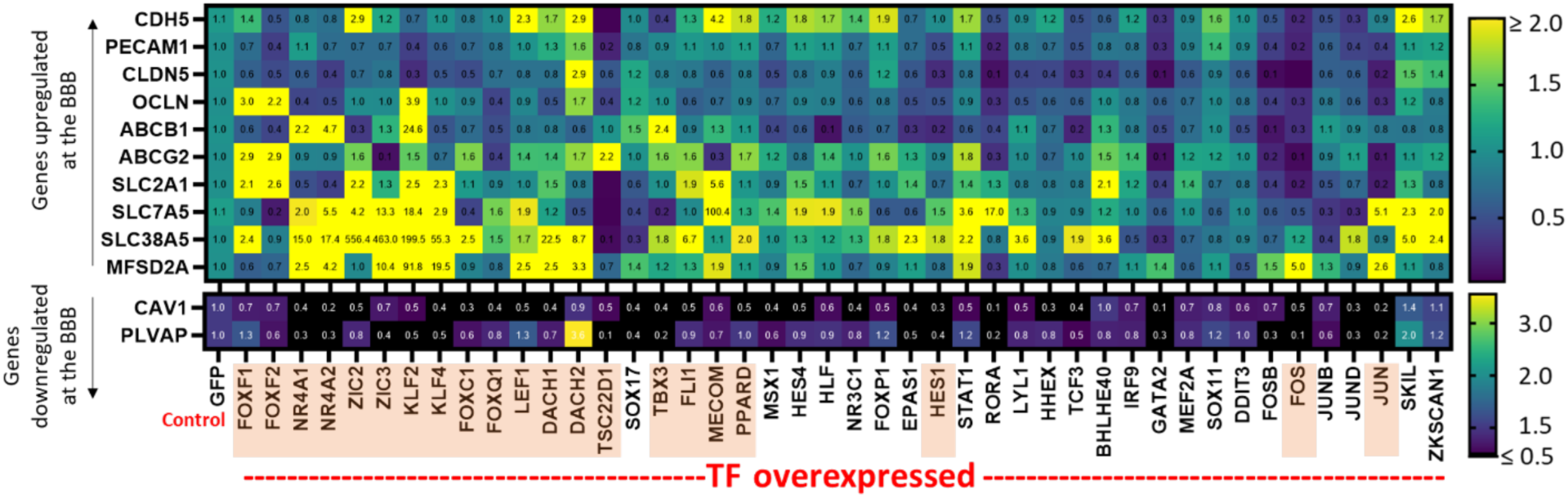
RT-qPCR screen of fpCECs with overexpression of a single TF. Forward programmed IMR90-4-derived CECs (fpCECs) were prepared by dosing lentiviruses delivering a single TF at MOI=2. RNA was extracted from the fpCECs four days after lentivirus transduction and RT-qPCR was performed on samples for 10 genes expressed at the BBB (*CDH5, PECAM1, CLDN5, OCLN, ABCB1, ABCG2, SLC2A1, SLC7A5, SLC38A5, MFSD2A*) and two genes downregulated at the BBB (*CAV1, PLVAP*). Fold-change of gene expression compared to the GFP control is displayed in each cell and also represented by color gradient. For overexpression, the color bar for upregulation is capped at 2-fold, and for downregulation is capped at 0.5-fold for better visualization. N=1 transduction for each TF was performed and the fold-change in gene expression was calculated. TFs that were selected for further analysis are highlighted in orange color.

**Fig. S2.**
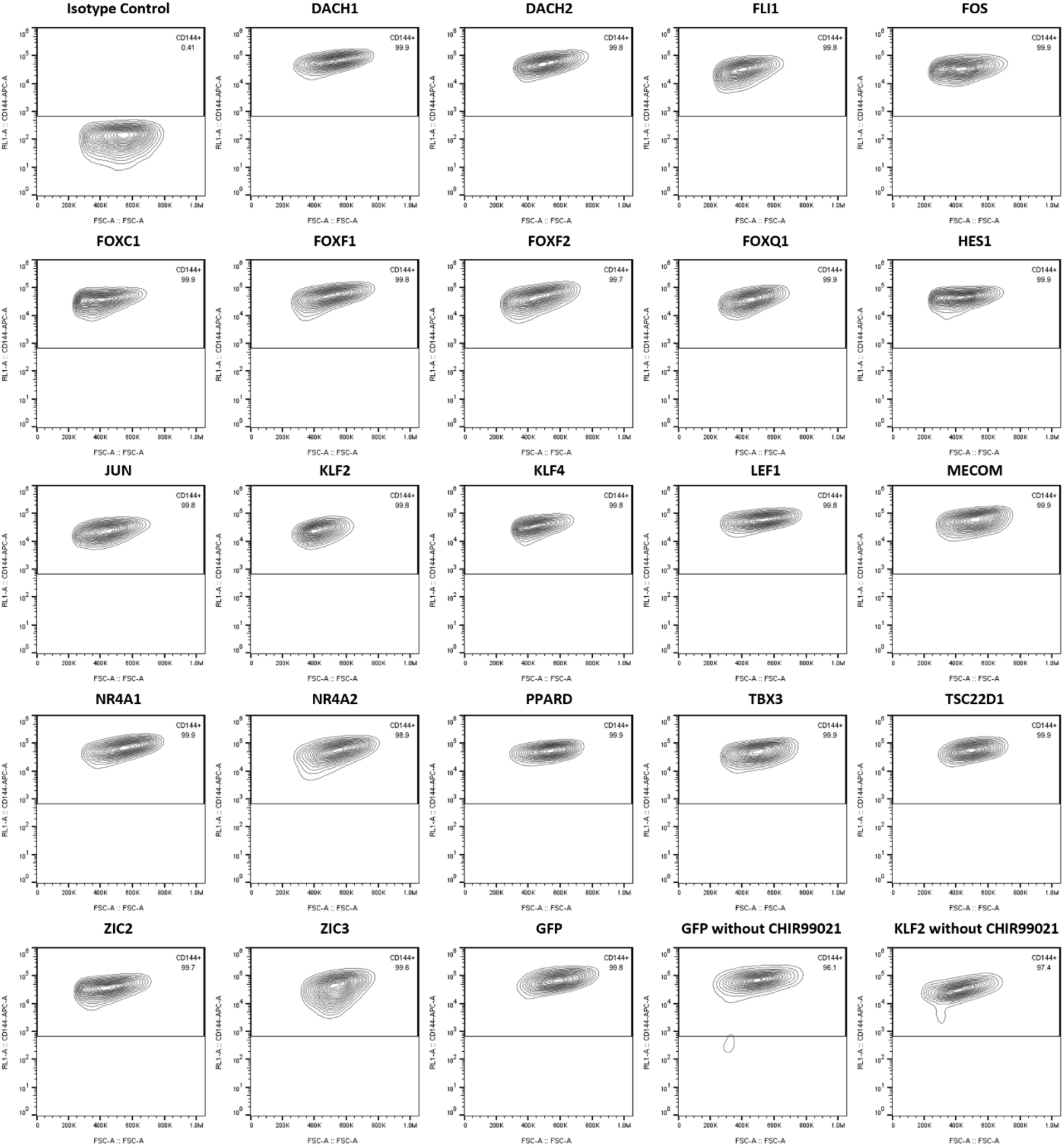
Endothelial marker VE-Cadherin (CDH5) quality check for bulk RNA-Seq samples. IMR90-4 hPSC-derived EPCs were dosed with lentivirus for GFP or TF overexpression at MOI=2 and simultaneously treated with CHIR99021 to activate Wnt signaling, unless otherwise specified. Cells were collected from a parallel transduced sample from the bulk RNA sequencing experiment for endothelial marker quality check. Flow cytometry plots show expression of endothelial cell marker CDH5 plotted against forward scatter area for all samples.

**Fig. S3.**
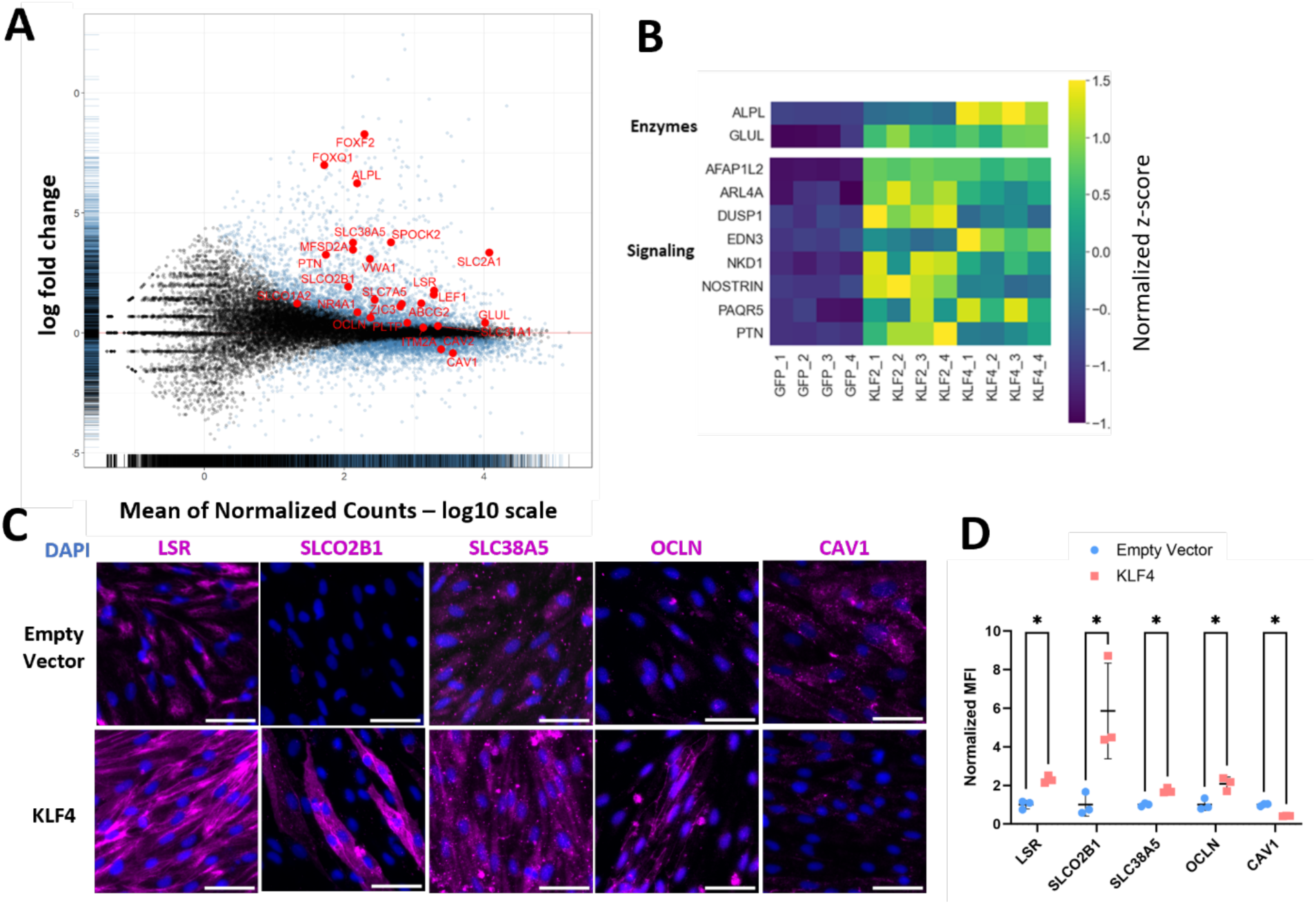
KLF family TFs induce BBB-like gene and protein expression. **(A)** MA plot for differential gene expression between fpCECs overexpressing *KLF4* and CECs overexpressing GFP. X-axis is the log_10_ mean of normalized counts in bulk RNA-seq. Y-axis is the log_2_(fold change) of gene expression. Some key BBB-enriched and depleted genes are marked in red. **(B)** Heatmap of the expression of BBB enzymes and signaling-related genes upregulated by *KLF2* and *KLF4* overexpression. z-scores normalized to the mean expression for each gene are plotted, with N=4 biological replicates per condition. fpCECs received either a control GFP lentivirus, *KLF2* lentivirus, or *KLF4* lentivirus at MOI=2. **(C)** Immunocytochemistry of specialized transporters (SLCO2B1, SLC38A5), tight junction (LSR, OCLN) and endocytosis-related marker (CAV1) in fpCECs receiving either an empty vector lentivirus or *KLF4* lentivirus at MOI=2. Scale bar is 50µm. **(D)** Quantification of immunocytochemistry images in panel D. N=3 biological replicates per condition. *:p<0.05 by Student’s t-test.

**Fig. S4.**
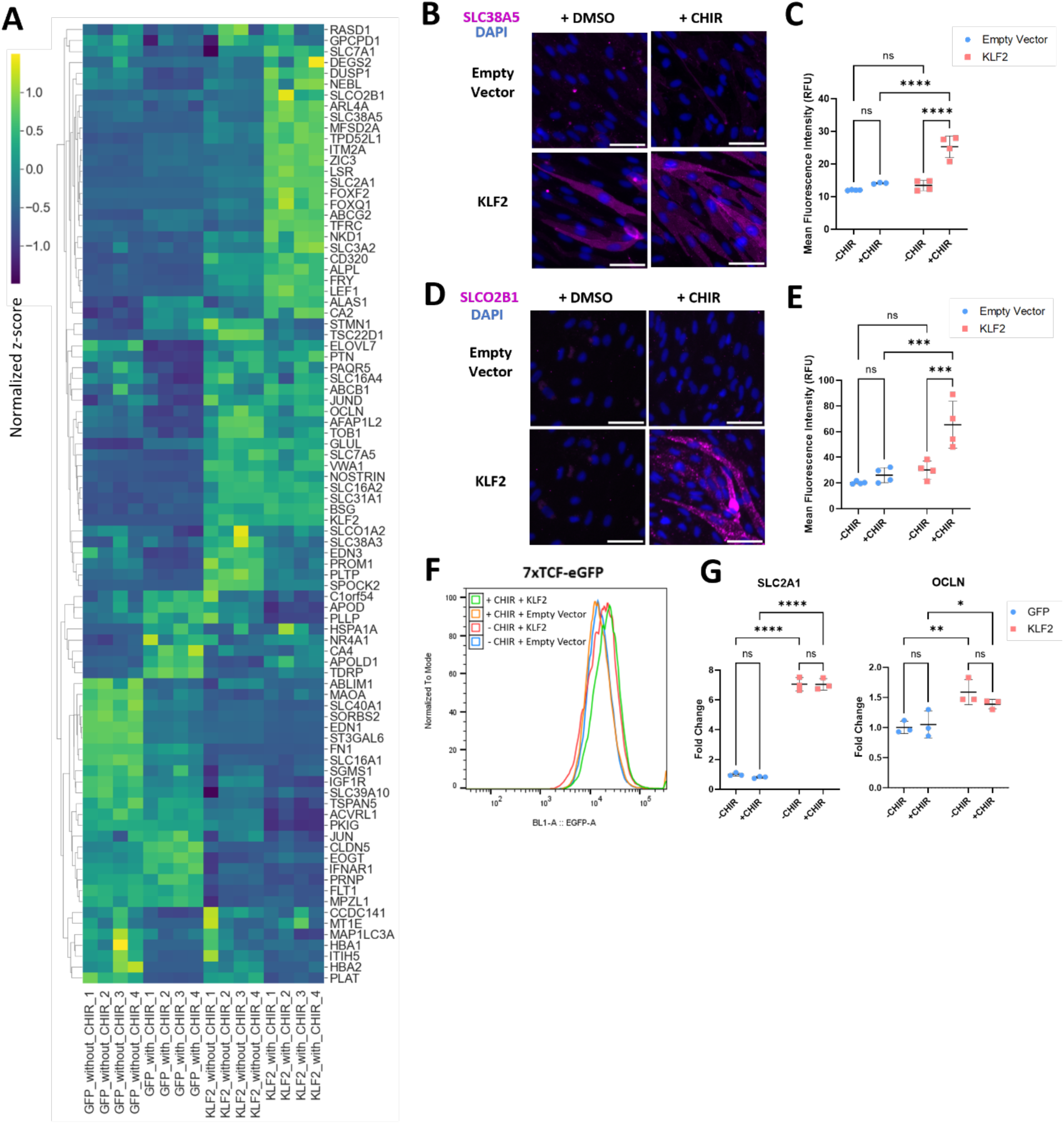
*KLF2* and Wnt activation synergistically induce BBB-like gene and protein expression. **(A)** Heatmap of the expression of 88 BBB-enriched genes in EPCs overexpressing *KLF2*. EPCs received either Wnt agonist CHIR99021 or DMSO control, and either *KLF2* lentivirus or GFP control lentivirus. N=4 biological replicates per condition are plotted. **(B)** Immunocytochemistry of neutral amino acid transporter SLC38A5 in EPCs receiving either Wnt agonist CHIR99021 or DMSO control, and either *KLF2* lentivirus or empty lentivirus. Scale bar is 50µm. **(C)** Quantification of SLC38A5 immunocytochemistry images. N=4 biological replicates per condition. ****:p<0.0001 in two-way ANOVA followed by Tukey’s post-hoc HSD test. **(D)** Immunocytochemistry of an organic anion transporter SLCO2B1 in EPCs receiving either Wnt agonist CHIR99021 or DMSO control, and either *KLF2* lentivirus or empty lentivirus. Scale bar is 50µm. **(E)** Quantification of SLCO2B1 immunocytochemistry images. N=4 biological replicates per condition. ***:p<0.001 in two-way ANOVA followed by Tukey’s post-hoc HSD test. **(F)** Representative flow cytometry histograms of eGFP expression using 7xTCF-eGFP H9 (7TGP) Wnt reporter line-derived EPCs transduced with either empty vector or *KLF2* lentiviruses, and with or without CHIR99021 treatment. Quantification in figure 3(O). **(G)** qPCR quantification of *SLC2A1* and *OCLN* gene expression in HUVECs transduced with GFP or *KLF2* lentivirus and cultured with or without CHIR99021. Expression is normalized to samples treated with empty vector lentivirus in the absence of CHIR99021. N=3 biological replicates. *:p<0.05; **:p<0.01; ***:p<0.001 in two-way ANOVA followed by Tukey’s post-hoc HSD test.

**Fig. S5.**
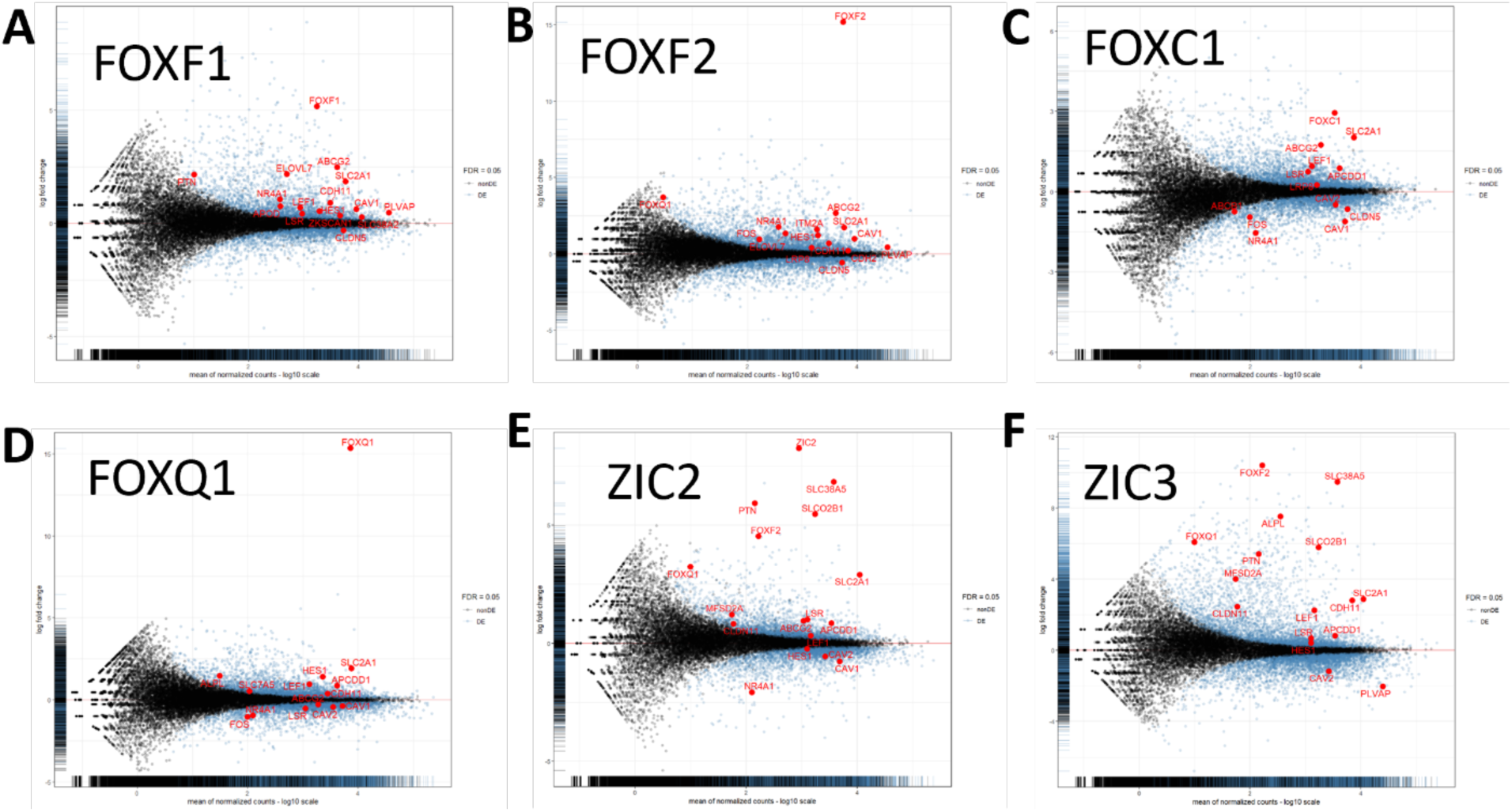
FOX and ZIC family TFs induce a subset of BBB-like gene expression changes. MA plots for differential gene expression between fpCECs overexpressing **(A)** *FOXF1*, **(B)** *FOXF2*, **(C)** *FOXC1*, **(D)** *FOXQ1*, **(E)** *ZIC2*, or **(F)** *ZIC3* and CECs overexpressing GFP. X-axis is the log_10_ (mean of normalized counts). Y-axis is the log_2_ (fold change of gene expression). Selected BBB-enriched and depleted genes are marked red.

**Fig. S6.**
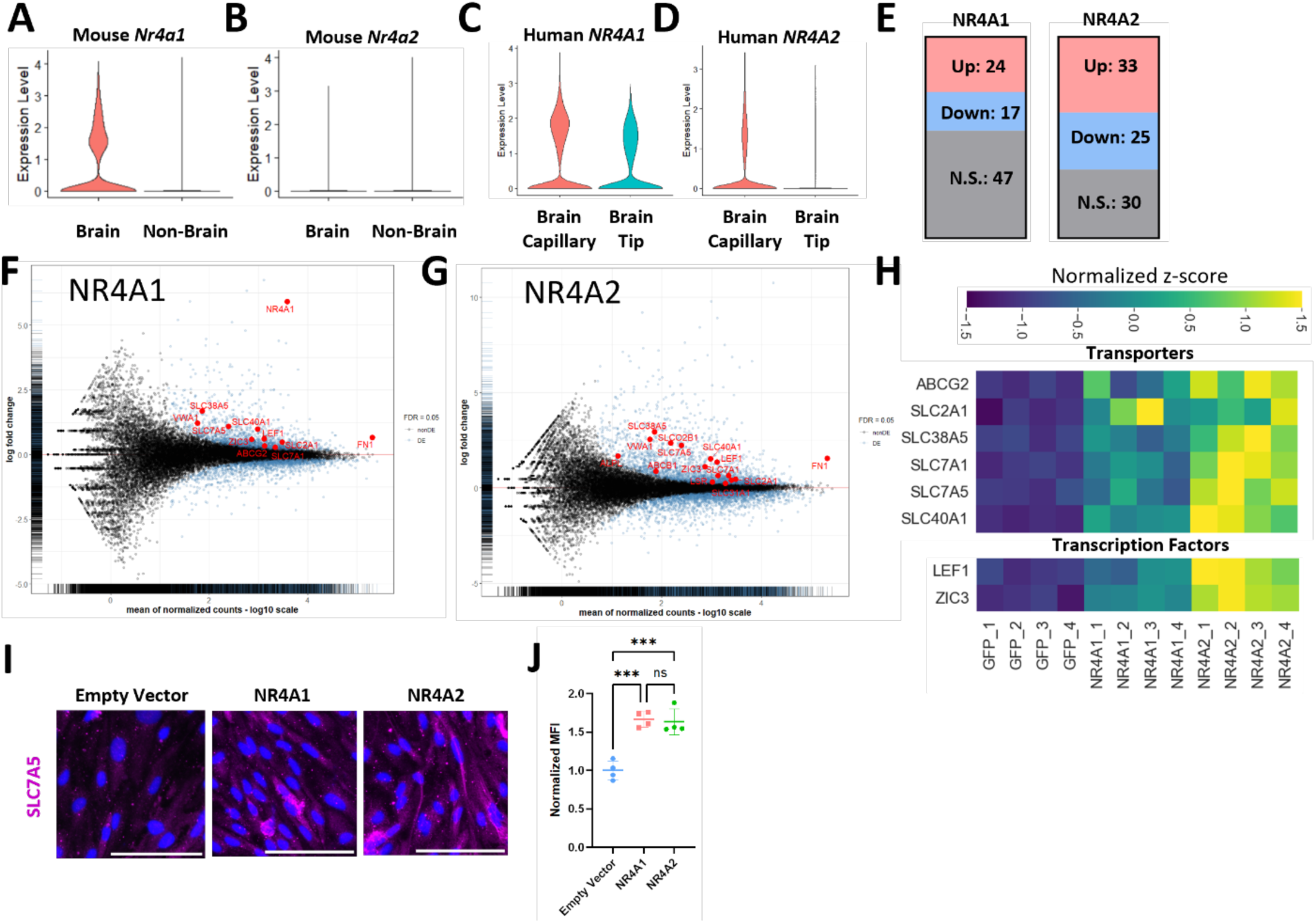
*NR4A1* and *NR4A2* TFs induce a subset of BBB-like gene expression changes. Expression of **(A)** *Nr4a1* and **(B)** *Nr4a2* in mouse brain and non-brain (heart, lung, liver, kidney and skeletal muscle) ECs in single cell transcriptomic analysis (*1*). Similarly, expression of **(C)** *NR4A1* and **(D)** *NR4A2* in human brain capillary and tip ECs was plotted (*2*). **(E)** Number of 88 BBB-enriched genes upregulated, downregulated, or exhibiting no significant changes upon overexpression of *NR4A1* or *NR4A2* in fpCECs. Adjusted p-value < 0.05 using the Wald test was used to determine whether upregulation or downregulation was statistically significant. **(F-G)** MA plot for differential gene expression between fpCECs overexpressing **(F)** *NR4A1* or **(G)** *NR4A2* and CECs overexpressing GFP. X-axis is the log_10_ (mean of normalized counts) in bulk RNA-seq. Y-axis is the log_2_ (fold change of gene expression). BBB-enriched and depleted genes are marked red. **(H)** Heatmap of the expression of genes regulating BBB transporters and transcription factor genes that are upregulated by *NR4A1* and *NR4A2* overexpression. Z-scores normalized to the mean expression for each of the BBB-enriched genes are plotted. N=4 biological replicates per condition. fpCECs receiving either a control GFP lentivirus, or TF lentivirus at MOI=2. **(I)** Immunocytochemistry of SLC7A5 in fpCECs receiving either an empty vector lentivirus, or *NR4A1* or *NR4A2* TF lentivirus at MOI=2. Scale bar is 100µm. **(J)** Quantification of SLC7A5 immunocytochemistry images. N=4 biological replicates per condition. MFI: mean fluorescence intensity. ***:p<0.001 in one-way ANOVA followed by Tukey’s post-hoc HSD test.

**Fig. S7.**
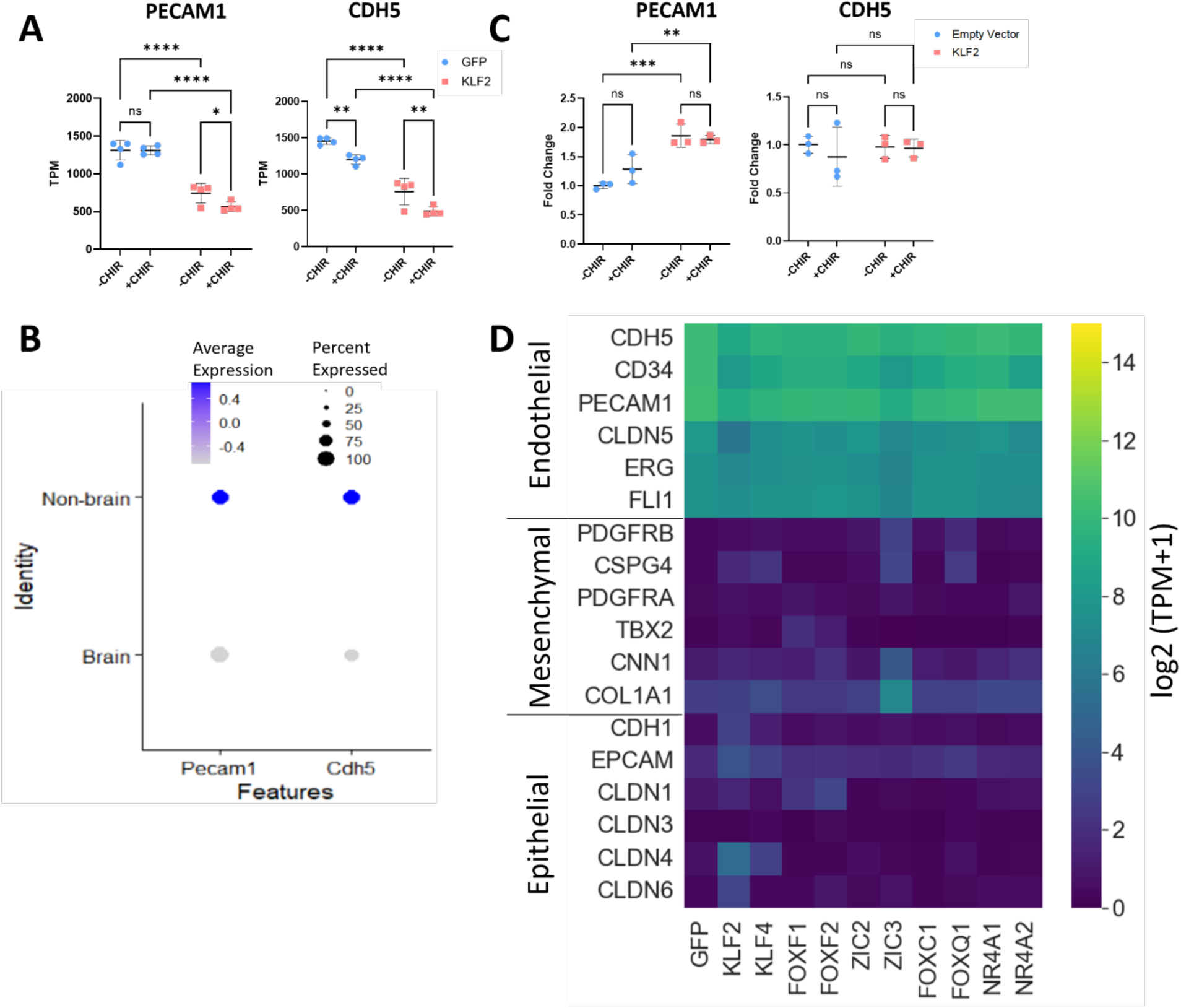
fpCECs with single TF overexpression maintain endothelial signature. **(A)** Quantification of *PECAM1* and *CDH5* gene expression in EPCs transduced with GFP or *KLF2* lentivirus and cultured with or without CHIR99021. *:p<0.05; **:p<0.01; ****:p<0.0001 in two-way ANOVA followed by Tukey’s post-hoc HSD test. **(B)** Dot plot for the expression of endothelial markers *Pecam1* and *Cdh5* in mouse scRNA-seq data for brain ECs and peripheral ECs (kidney, lung, heart, liver and skeletal muscle) (*1*). **(C)** qPCR quantification of *PECAM1* and *CDH5* gene expression in HUVECs transduced with GFP or *KLF2* lentivirus and cultured with or without CHIR99021. Expression is normalized to samples treated with empty vector lentivirus in the absence of CHIR99021. N=3 biological replicates. *:p<0.05; **:p<0.01; ***:p<0.001 in two-way ANOVA followed by Tukey’s post-hoc HSD test. **(D)** Heatmap of the expression of endothelial, mesenchymal, and epithelial transcripts in CECs receiving either *KLF2, KLF4, FOXF1, FOXF2, ZIC2, ZIC3, FOXC1, FOXQ1, NR4A1, NR4A2* lentiviruses or GFP control lentivirus. Log_2_ (TPM+1) was plotted.

**Fig. S8.**
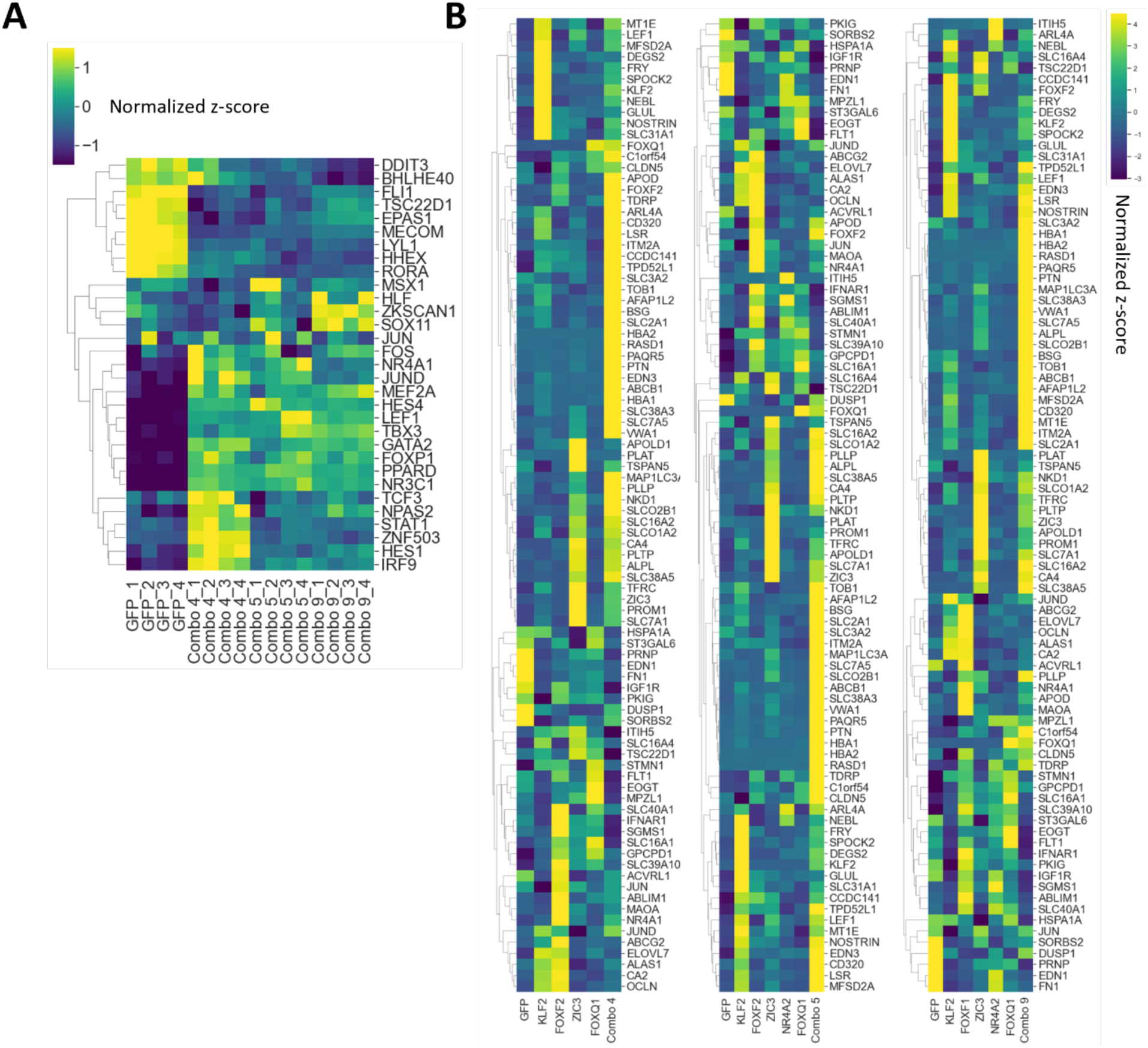
Combinatorial TF overexpression reveals synergy among TFs to induce BBB-like gene expression. **(A)** Heatmap of the expression of BBB-enriched TFs from Figure 1 in fpCECs overexpressing either GFP or TF combos with the highest BBB scores (Combos 4, 5 and 9). BBB TFs enriched in at least two out of the three scRNA-seq datasets shown in Figure 1 are presented in this panel. Those already part of the TF combos are excluded from this panel. Average z-score normalized to the mean expression for each gene for N=4 biological replicates is plotted. **(B)** Heatmap of the expression of 88 BBB-enriched genes in fpCECs overexpressing TF combos 4, 5, and 9, along with the single TFs included in the combinations. Average z-score normalized to the mean expression for each gene for N=4 biological replicates is plotted.

**Fig. S9.**
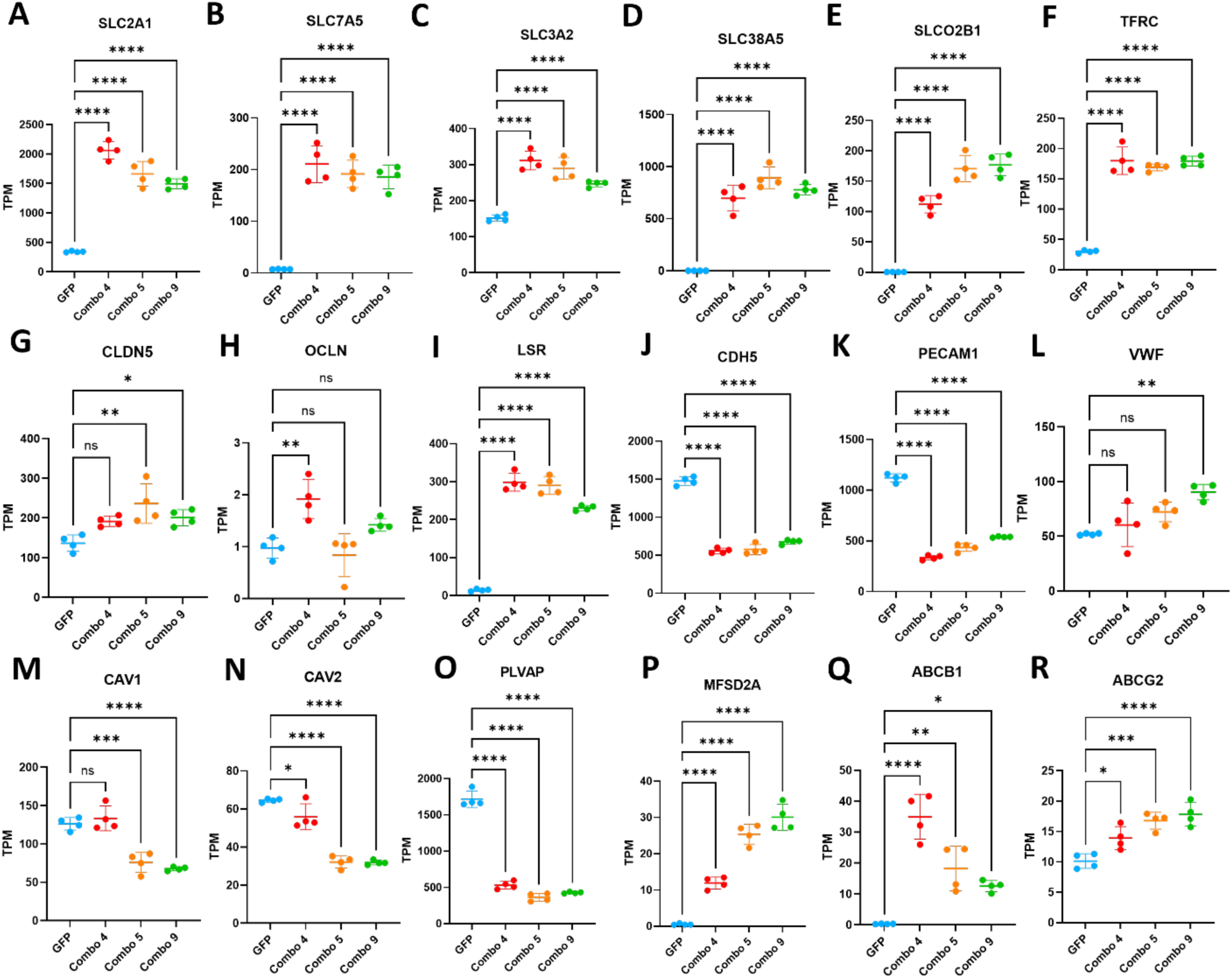
Expression levels of selected BBB genes in fpCECs. Transcript per million (TPM) values of selected transcripts for fpCECs transduced with GFP control lentivirus or TF Combo 4, 5, or 9 lentiviruses. The top row displays expression of **(A)** glucose transporter (*SLC2A1*), **(B-E)** other solute carriers enriched in brain ECs (*SLC7A5, SLC3A2, SLC38A5, SLCO2B1*), and the **(F)** transferrin receptor gene (*TFRC*). The second row displays expression of **(G-I)** junctional markers (*CLDN5, OCLN, LSR*), and **(J-L)** endothelial cell markers (*CDH5, PECAM1, VWF*). The third row displays **(M-P)** transcytosis related markers (*CAV1, CAV2, PLVAP, MFSD2A*) and **(Q-R)** efflux transporters (*ABCB1, ABCG2*). TPM data for all genes are in supplementary file 1. *:p<0.05, **:p<0.01, ***:p<0.001, ****: p<0.0001 in one-way ANOVA followed by Dunnett’s post-hoc test.

**Fig. S10.**
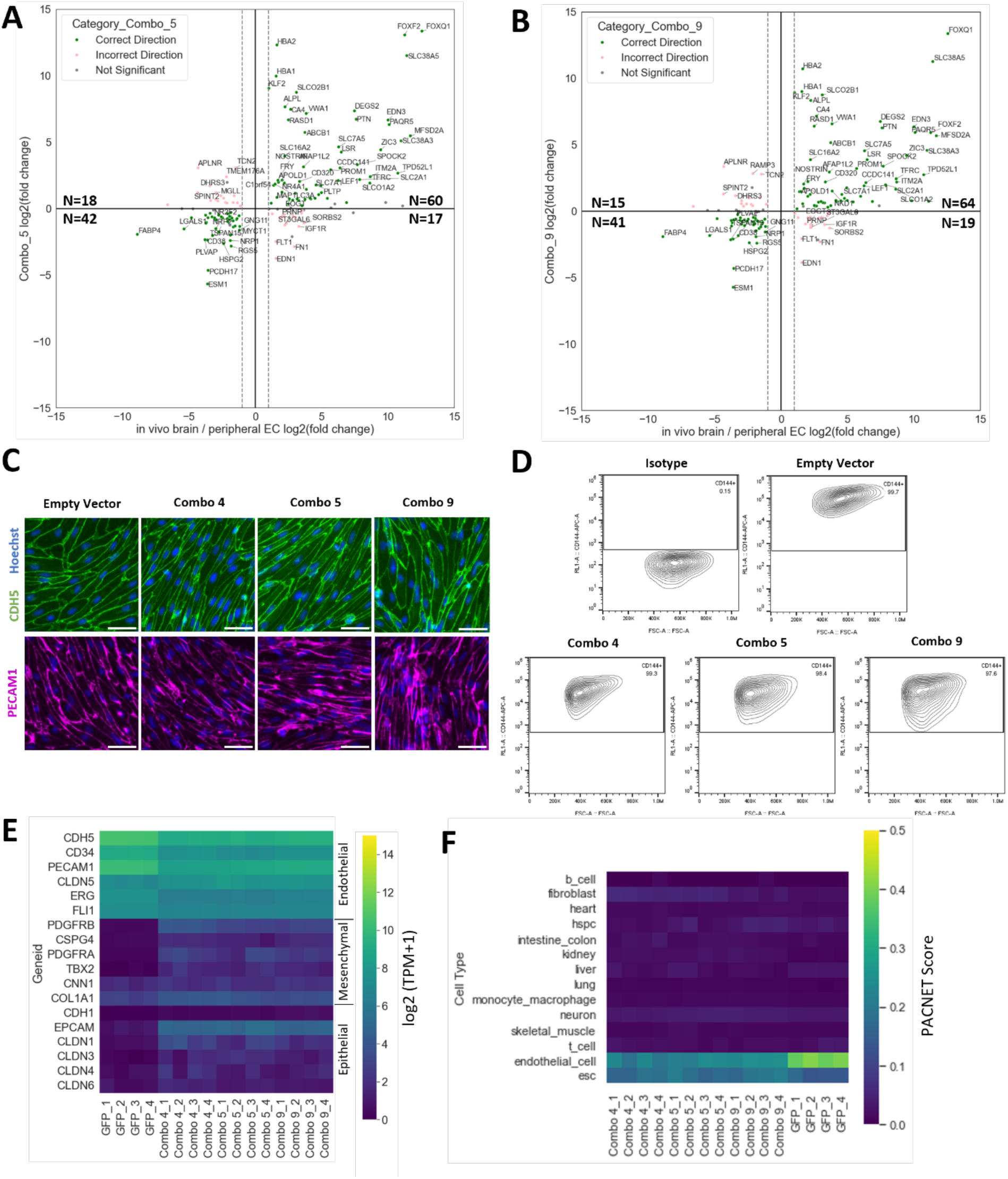
Combinatorial TF overexpression induces BBB properties in fpCECs while maintaining endothelial identity. Scatter plot comparing gene expression changes elicited by. **(A)** Combo 5 TF, or **(B)** Combo 9 TF overexpression to gene expression difference between brain and non-brain ECs *in vivo*. Plotting strategy is the same as Figure 6B. Green and pink dots represent genes significantly changing in the correct and incorrect direction, respectively, padj<0.05. Gray dots represent non-significant genes. Number of significant genes in each quadrant are annotated. **(C)** Immunocytochemistry analysis of endothelial markers CDH5 and PECAM1 in fpCECs treated with either empty vector lentiviruses or Combo 4, 5, or 9 lentiviruses. Scale bar is 50µm. **(D)** Flow cytometry analysis of fpCECs with empty vector control or TF combos 4, 5, or 9 lentivirus treatment for expression of endothelial cell marker CDH5. **(E)** Heatmap of the expression of endothelial, mesenchymal and epithelial transcripts in fpCECs. Samples are fpCECs receiving Combo 4, 5, or 9 TF lentiviruses or GFP control lentivirus. N=4 biological replicates per condition are plotted. **(F)** PACNet analysis of cell fate for fpCECs overexpressing GFP or Combo 4, 5, or 9 TFs. PACNet scores for cell types are shown.

**Fig. S11.**
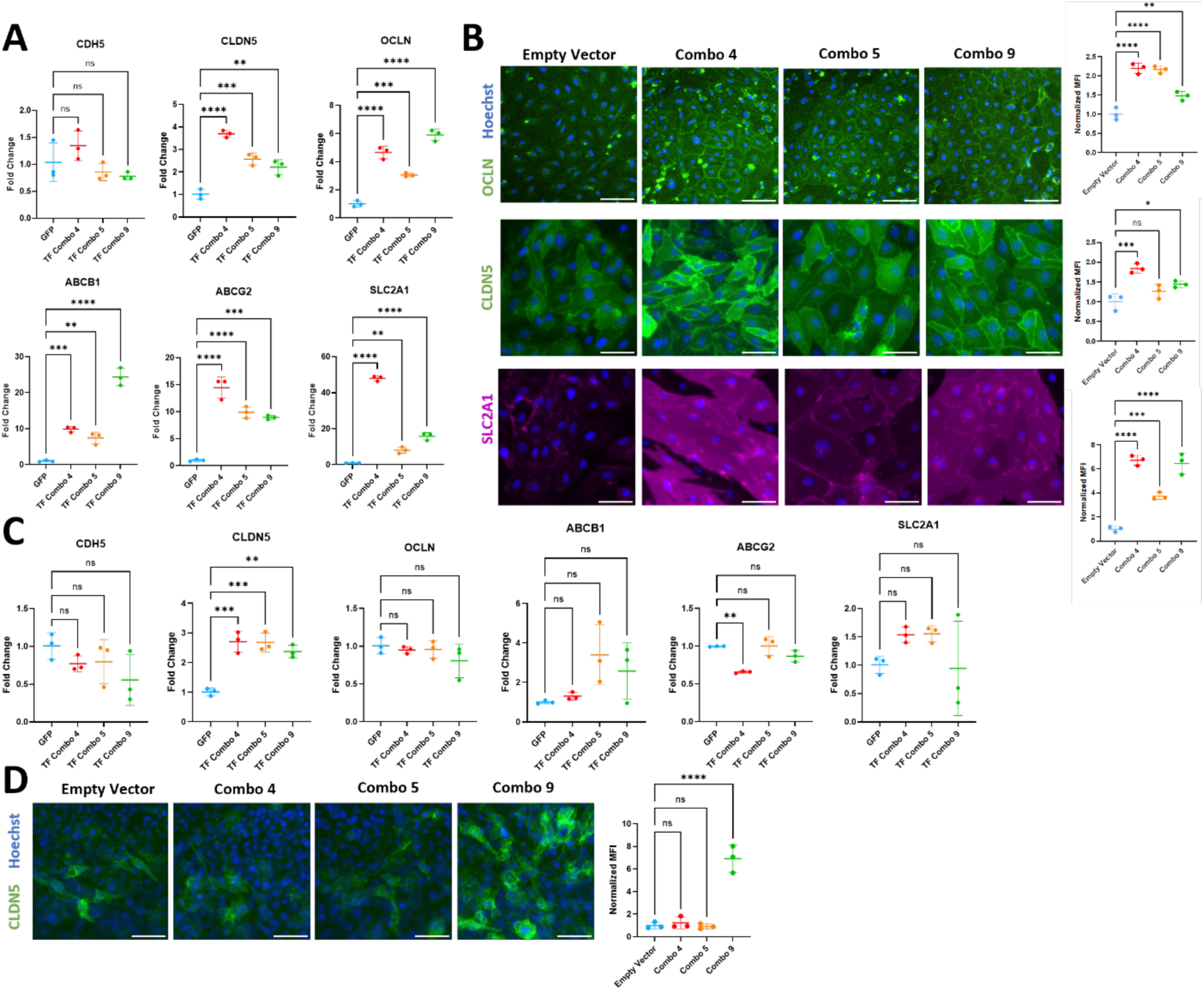
Combinatorial TF overexpression induces HUVECs and hCMEC/D3 cells to gain BBB-like gene and protein expression. **(A)** HUVECs were dosed with TF Combo 4, 5, and 9 lentiviruses at MOI=2 for each TF or GFP control. RNA was extracted four days after lentivirus transduction and RT-qPCR was performed on samples for indicated genes. Fold changes of *CDH5*, *CLDN5*, *OCLN*, *ABCB1*, *ABCG2*, *SLC2A1* gene expression normalized to the GFP control are plotted. One-way ANOVA followed by Dunnett’s post-hoc test was used for statistical analysis. N=3 biological replicates. **: p<0.01, ***: p<0.001; ****:p<0.0001. **(B)** Immunocytochemistry of OCLN, CLDN5 and SLC2A1 in HUVECs receiving either an empty vector lentivirus, or TF combo 4, 5 or 9 lentiviruses. Scale bar is 50µm. MFI of N=3 biological replicates was quantified for each condition. One-way ANOVA followed by Dunnett’s post-hoc test was used for statistical analysis. *:p<0.05; **: p<0.01, ***: p<0.001; ****:p<0.0001. **(C)** hCMEC/D3 cells were dosed with GFP or TF Combo 4, 5, and 9 lentiviruses for each condition. RNA was extracted four days after lentivirus transduction and RT-qPCR was performed on samples for indicated genes. Fold changes of *CDH5*, *CLDN5*, *OCLN*, *ABCB1*, *ABCG2*, *SLC2A1* gene expression normalized to the GFP control are plotted. One-way ANOVA followed by Dunnett’s post-hoc test was used for statistical analysis. N=3 biological replicates. **: p<0.01, ***: p<0.001. **(D)** Immunocytochemistry of CLDN5 in hCMEC/D3 cells receiving either an empty vector lentivirus, or TF combo 4, 5, or 9 lentiviruses. MFI of N=3 biological replicates was quantified for each condition. One-way ANOVA followed by Dunnett’s post-hoc test was used for statistical analysis. ****: p<0.0001.

**Fig. S12.**
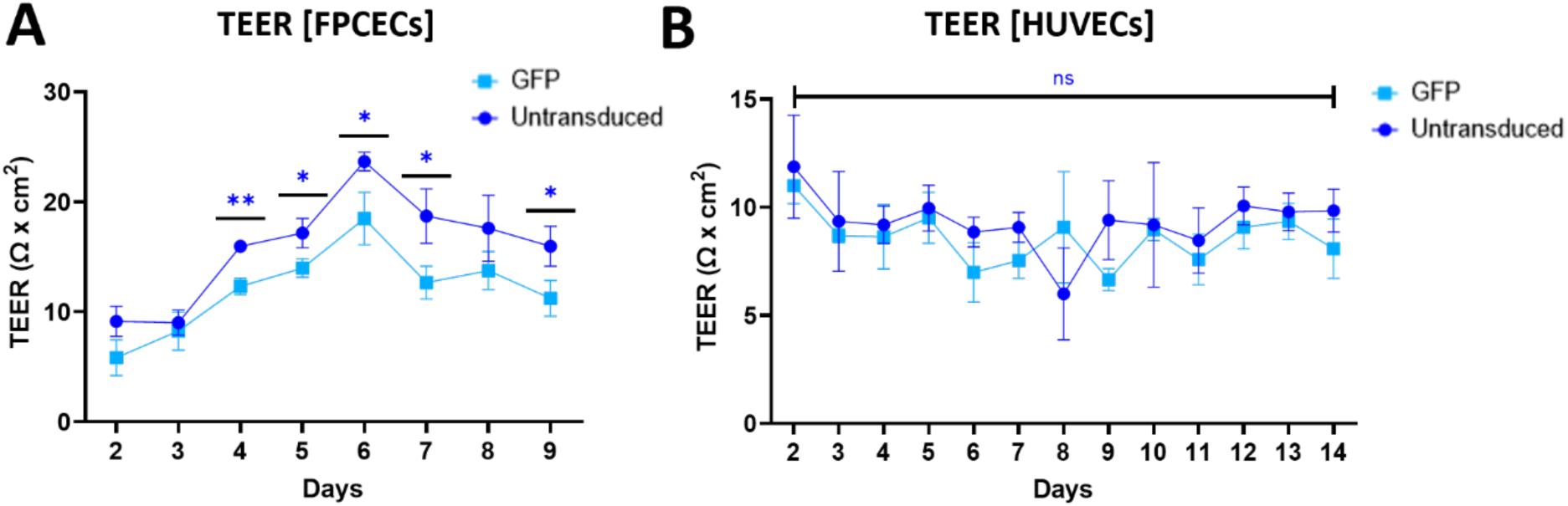
Effects of lentivirus treatment on CECs and HUVECs TEER. TEER values were measured for. **(A)** fpCECs and **(B)** HUVECs that were transduced with GFP control lentivirus on Day 1 or left untransduced. TEER measurements began on Day 2. Mean ± SD of N=3 biological replicates is plotted. ns: non-significant, *:p<0.05, **:p<0.01 in an unpaired t-test.

**Table S1.**
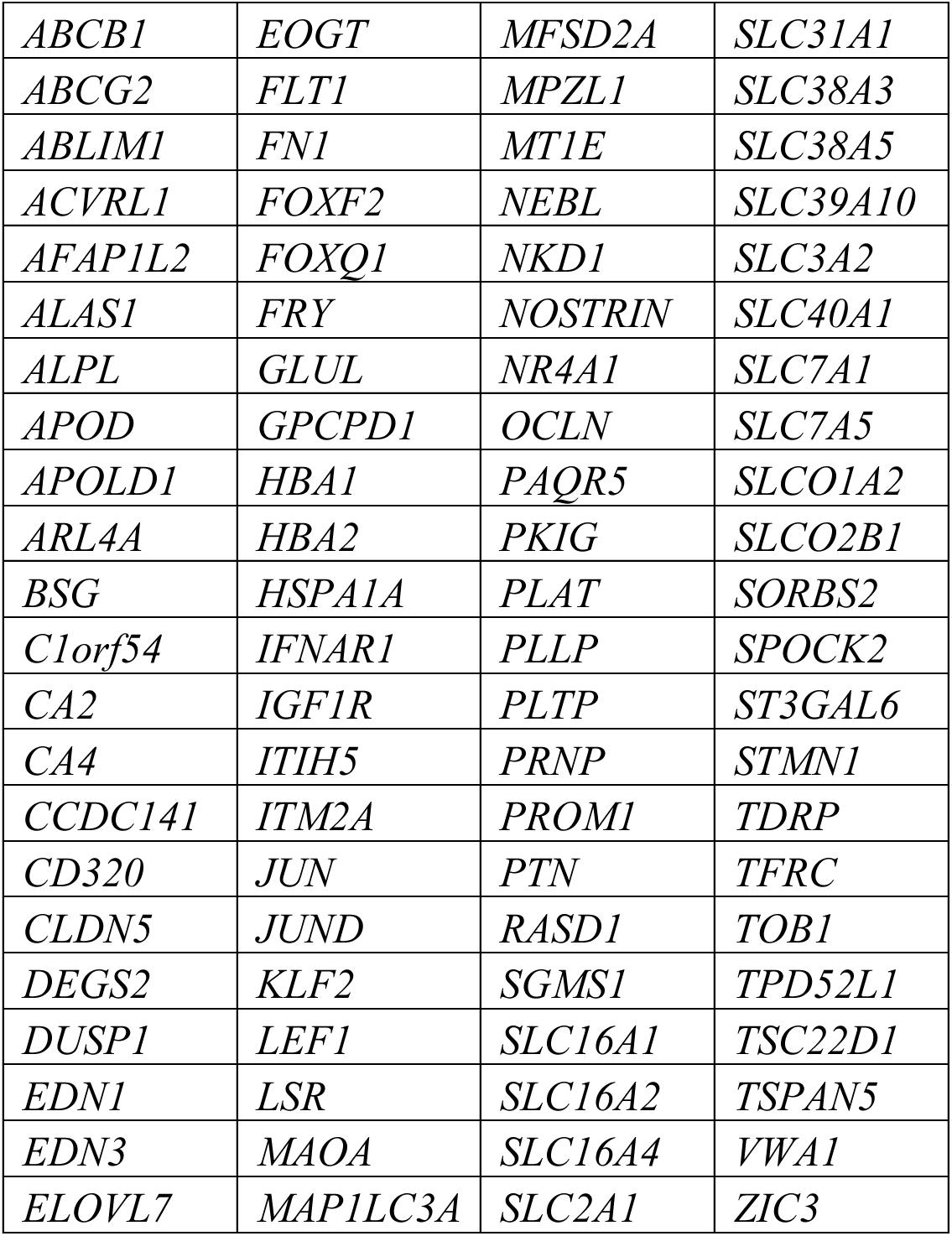
Panel of BBB-enriched genes mined through scRNA-seq analysis.

**Table S2.** Addgene plasmids for GFP control and TFs used in this study.

| **Addgene Catalog Number** | **TFORF Number** | **Gene/Insert Name** |
| --- | --- | --- |
| 145025 | TFORF3549 | *GFP* |
| 143370 | TFORF2408 | *FOXF1* |
| 144021 | TFORF2409 | *FOXF2* |
| 144416 | TFORF2940 | *NR4A1* |
| 142874 | TFORF1856 | *NR4A2* |
| 143984 | TFORF1884 | *ZIC2* |
| 141739 | TFORF0859 | *ZIC3* |
| 142715 | TFORF0987 | *KLF2* |
| 142249 | TFORF0985 | *KLF4* |
| 142534 | TFORF0102 | *FOXC1* |
| 142528 | TFORF0024 | *FOXQ1* |
| 141862 | TFORF1659 | *LEF1* |
| 144161 | TFORF2901 | *DACH1* |
| 143660 | TFORF2905 | *DACH2* |
| 141786 | TFORF1040 | *TSC22D1* |
| 141998 | TFORF2260 | *SOX17* |
| 142519 | TFORF1973 | *TBX3* |
| 141911 | TFORF2079 | *FLI1* |
| 143395 | TFORF0337 | *MECOM* |
| 143862 | TFORF1967 | *PPARD* |
| 141692 | TFORF0769 | *MSX1* |
| 143583 | TFORF2286 | *HES4* |
| 141513 | TFORF0099 | *HLF* |
| 143497 | TFORF1432 | *NR3C1* |
| 142485 | TFORF0213 | *FOXP1* |
| 143762 | TFORF1106 | *EPAS1* |
| 142008 | TFORF2283 | *HES1* |
| 144562 | TFORF3086 | *STAT1* |
| 144463 | TFORF2987 | *RORA* |
| 143126 | TFORF1497 | *LYL1* |
| 142805 | TFORF1496 | *HHEX* |
| 142580 | TFORF0273 | *TCF3* |
| 142336 | TFORF2593 | *BHLHE40* |
| 144933 | TFORF3457 | *IRF9* |
| 144474 | TFORF2998 | *GATA2* |
| 144489 | TFORF3013 | *MEF2A* |
| 142927 | TFORF2258 | *SOX11* |
| 144595 | TFORF3119 | *DDIT3* |
| 144244 | TFORF2815 | *FOSB* |
| 144413 | TFORF2937 | *FOS* |
| 144709 | TFORF3233 | *JUNB* |
| 141685 | TFORF0756 | *JUND* |
| 142292 | TFORF1932 | *JUN* |
| 142619 | TFORF0421 | *SKIL* |
| 141919 | TFORF2096 | *ZKSCAN1* |

**Table S3.**
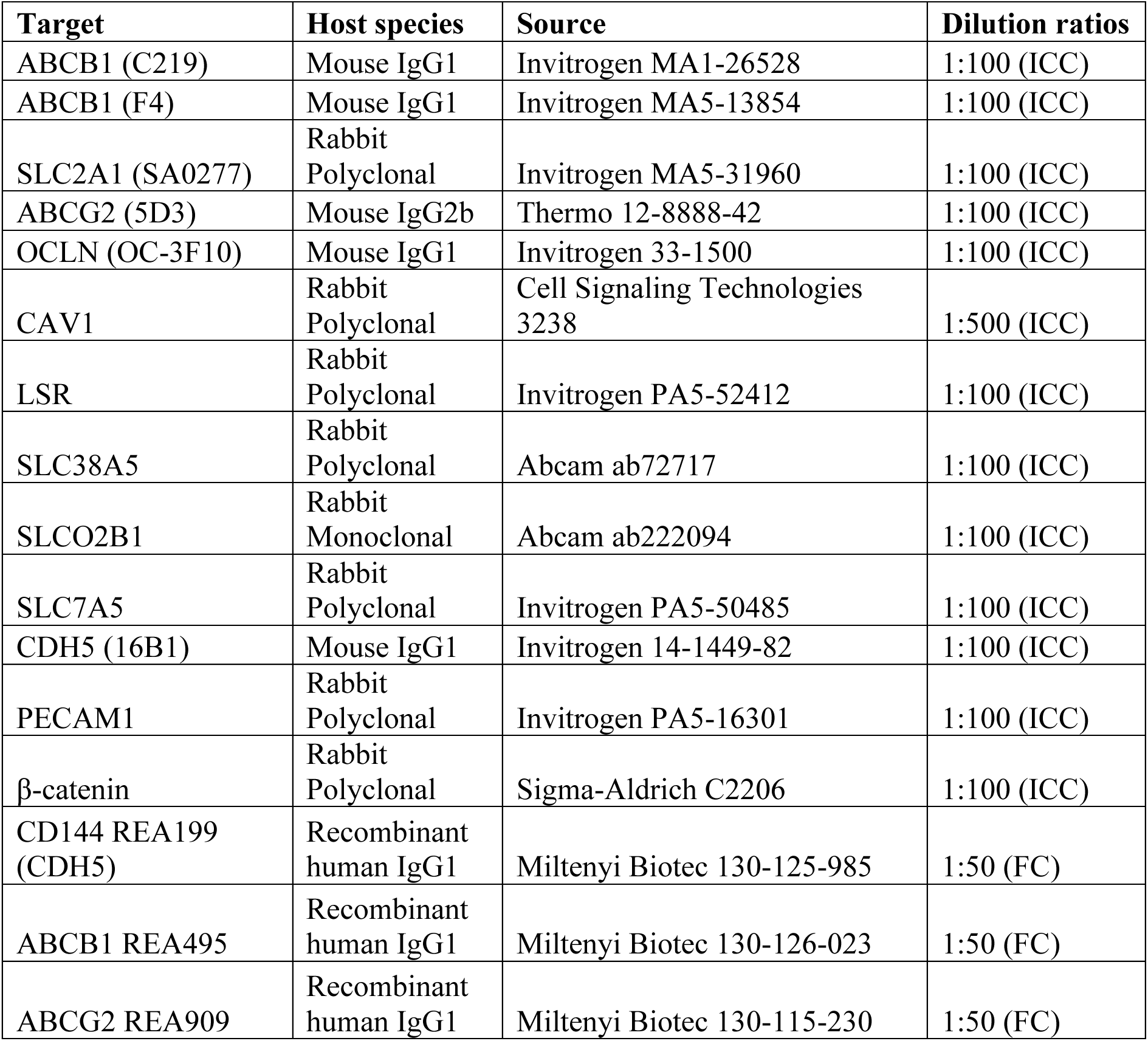
Antibodies used in this study.

**Table S4.**
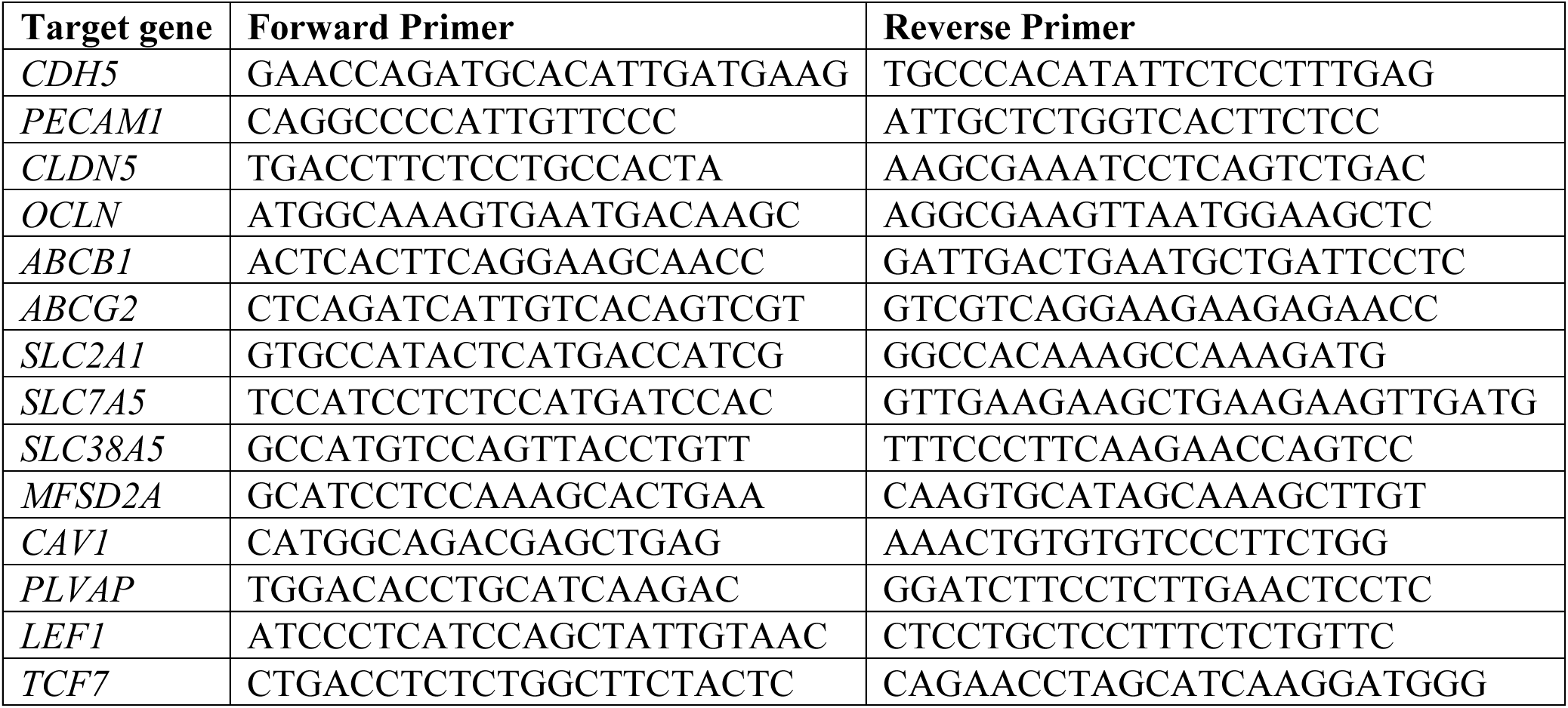
Primer sequences for RT-qPCR used in this study.

## Descriptions of Files S1 to S5 (separate files)

**File S1**.

TPM values of all samples (single TFs and combinations).

**File S2**.

Absolute values, fold changes, or statistics corresponding to heatmaps in figures.

**File S3**.

TF combinations prediction scores.

**File S4**.

List of genes enriched and depleted in brain ECs.

**File S5**.

BBB gene expression changes induced by TF combinations and the direction of the change.

